# Ethnicity-specific transcriptomic variation in immune cells and correlation with disease activity in systemic lupus erythematosus

**DOI:** 10.1101/2020.10.30.362715

**Authors:** Gaia Andreoletti, Cristina M. Lanata, Ishan Paranjpe, Tia S. Jain, Joanne Nititham, Kimberly E. Taylor, Alexis J Combes, Lenka Maliskova, Chun Jimmie Ye, Patricia Katz, Maria Dall’Era, Jinoos Yazdany, Lindsey A. Criswell, Marina Sirota

## Abstract

Systemic lupus erythematosus (SLE) is a heterogeneous autoimmune disease in which outcomes vary among different racial groups. The aim of this study is to leverage large-scale transcriptomic data from diverse populations to better sub-classify SLE patients into more clinically actionable groups. We leverage cell sorted RNA-seq data (CD14^+^ monocytes, B cells, CD4^+^T cells, and NK cells) from 120 SLE patients (63 Asian and 57 White individuals) and apply a four tier analytical approach to identify SLE subgroups within this multiethnic cohort: unsupervised clustering, differential expression analyses, gene co-expression analyses, and machine learning. K-means clustering on the individual cell type data resulted in three clusters for CD4 and CD14, and two clusters for B cells and NK cells. Correlation analysis revealed significant positive associations between the transcriptomic clusters of each immune cell and clinical parameters including disease activity and ethnicity. We then explored differentially expressed genes between Asian and White groups for each cell-type. The shared differentially expressed genes across the four cell types were involved in SLE or other autoimmune related pathways. Co-expression analysis identified similarly regulated genes across samples and grouped these genes into modules. Samples were grouped into White-high, Asians-high (high disease activity defined by SLEDAI score >=6) and White-low, Asians-low (SLEDAI < 6). Random forest classification of disease activity in the White and Asian cohorts showed the best classification in CD4^+^ T cells in White. The results from these analyses will help stratify patients based on their gene expression signatures to enable precision medicine for SLE.

## Introduction

Systemic lupus erythematosus (SLE) is a complex multisystem autoimmune disorder characterized by dysregulation of the innate and adaptive arms of the immune system^1^. The predisposition and clinical phenotype of SLE, a disease associated with significant morbidity and mortality, are attributed to a combination of genetics, hormones, and environmental factors^2^. Heterogenous clinical and serologic manifestations, a waxing and waning course, and delays in diagnosis contribute to the complexity of this disease.

Genome-wide association studies have discovered ~100 susceptibility loci for SLE^3^. Many of these loci are unique to European-Americans patients while others have been identified for African American or Hispanic-Americans^4,5^. Susceptibility to SLE has a strong genetic component, and trans-ancestral genetic studies have revealed a substantial commonality of shared genetic risk variants across different genetic ancestries that predispose to the development of SLE^6^. More than half of patients with SLE show a dysregulation in the expression of genes in the interferon (IFN) pathway^7^. Scientific evidences have identified that the dominant group most commonly diagnosed with SLE are women of minority status in low socio-economic environments^8^. A recent epidemiologic study comparing lupus manifestations among four major racial and ethnic groups found substantial differences in the prevalence of several clinical SLE manifestations among racial/ethnic groups and discovered that African Americans, Asian/Pacific Islanders, and Hispanic patients are at increased risk of developing several severe manifestations following a diagnosis of SLE^9^. It is believed that an increased genetic risk burden in these populations, associated with increased autoantibody reactivity in non-white individuals with SLE, may explain the more severe lupus phenotype^6^. As patients with similar Systemic Lupus Erythematosus Disease Activity Index, or SLEDAI^10^, scores may have different prognoses and treatment responses^11^, there is an urgent need to establish a new method of stratification of lupus patients.

With the recent advances in molecular measurements and computational technologies, there are incredible opportunities to characterize disease associated genes and pathways. Previous studies have used machine learning (ML) and clustering approaches to try to stratify patients with SLE based on different parameters including clinical data, expression quantitative trait loci (eQTLs), methylation, and transcriptomic data^12–15^. However, these studies have been mainly conducted on whole blood or peripheral blood mononuclear cells (PBMCs), therefore not considering the involvement of different immune cell types in disease, and in White cohorts, thus without taking into account a relationship between disease activity and ethnic background.

In our own previous work, unsupervised clustering of the 18 American College of Rheumatology (ACR) classification criteria on an ethnically diverse lupus cohort revealed three stable clusters, characterized by significant differences in several SLE features as well as the lupus severity index^12^. Following up on the clinical clustering study, the objective of the current study which leverages the same cohort is to use large-scale transcriptomic data from diverse ethnic population to identify transcriptomic signature in SLE relating various clinical and demographic factors and better sub-classify SLE patients into more clinically actionable groups. More specifically, we leverage bulk RNA sequencing data on four immune-cell types sorted from peripheral blood mononuclear cells (PBMCs) (CD14^+^ monocytes, B cells, CD4^+^T cells and NK cells) of 120 patients (63 Asian and 57 White individuals) from a multi-racial/ethnic cohort of individuals with physician-confirmed SLE. This study aims to look for specific-transcriptomic effects in each of these immune cell types using a four-tier approach: unsupervised clustering, differential expression analyses, gene co-expression analyses, and machine learning approaches (Figure 1).

Our approach helped to identify patient sub-clusters based on their gene expression data and to describe the involvement of CD14^+^ monocytes, B cells, CD4^+^ T cells and NK cells in disease in a SLE multiethnic/racial cohort. This new classification might provide insightful information to improve predictions of disease outcomes and insight into subtype-specific mechanistic pathways that could be strategically targeted. We further explore molecular pathways that underlie the clinical and demographic differences in SLE.

## Results

### Unsupervised K-means clustering identified specific patient clusters per immune cell-type

After profiling 120 SLE patients for cell sorted bulk RNA-seq data (CD4^+^ T cells, CD14^+^ monocytes, B cells, and NK cells) from a multi-racial/ethnic cohort, a total of 10% of the samples were removed after QC filtering retaining 415 samples (91 NK, 105 B cells, 108 CD4^+^ T-cells, and 111 CD14^+^monocytes) (Figure S1A). Batch effects were taken into account in all the analyses using the limma software. Using the QC’ed data, we wanted to look for specific-transcriptomic effects in each of these cell types using a four-tier approach which included: unsupervised clustering; differential expression analyses, gene co-expression analyses, and random forest (Figure 1).

Unsupervised K-means clustering on the individual cell types was utilized to identify patient sub clusters. Clusters with a Jaccard mean stability score greater than 0.6 were considered stable and therefore only clusters with a score greater than or equal to 0.6 were retained for further analyses (Table S3). Clustering on CD4^+^T cells and CD14^+^ monocytes yielded three distinct clusters, while only 2 clusters were identified for B cells and NK cells (Figure 2A). Thus, we sought to test for association between the transcriptomic K-means clustering and the clinical and demographic parameters (Figure 2B).

**Figure 2:**
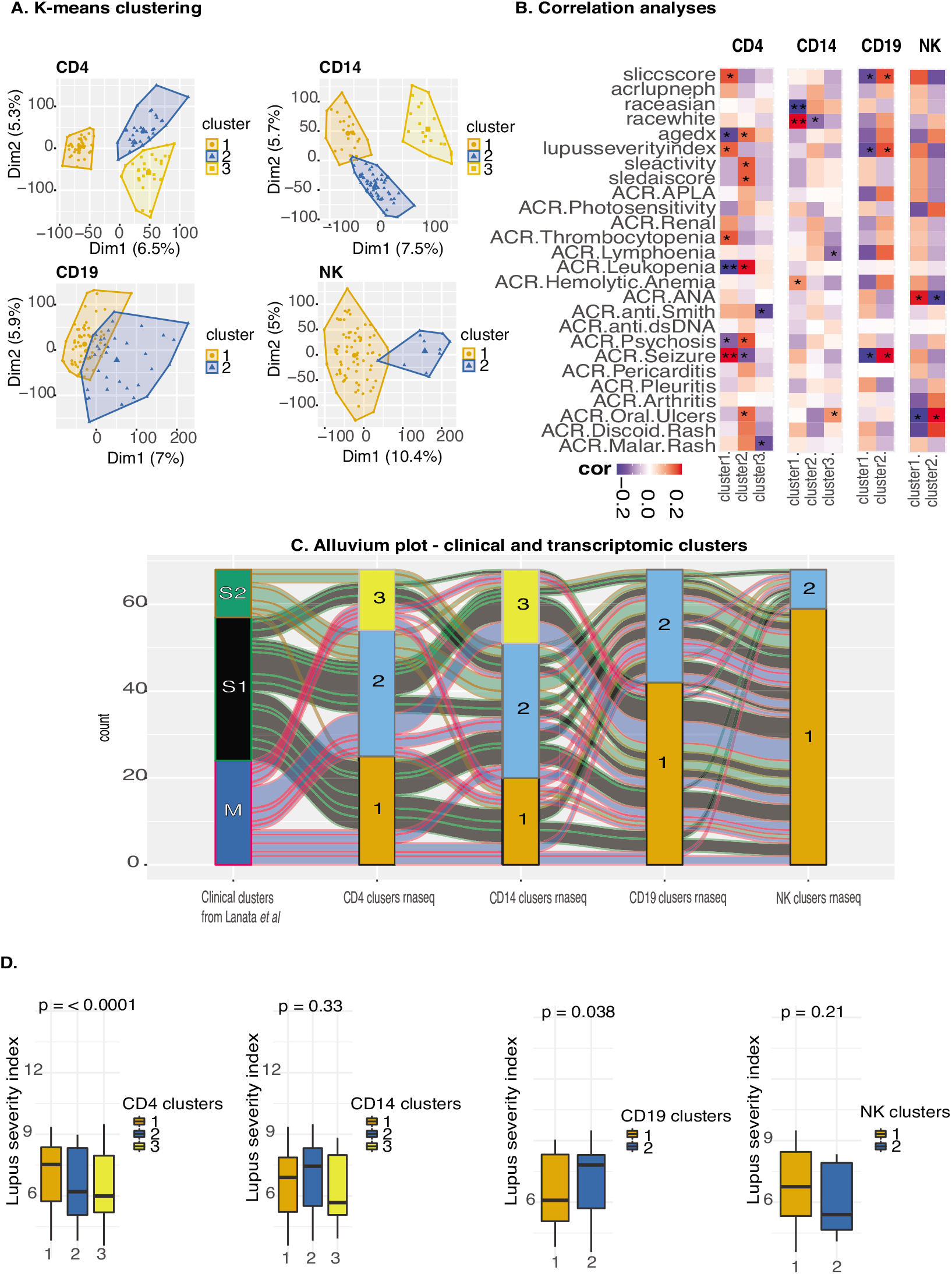
A) K-means clustering on the CD14^+^ monocytes, CD4^+^ T-cells, B cells and NK cells. B) association analyses between clinical parameters and K-means s clustering. Red – positive association, blue – negative association. Number of stars indicate the lever of significance. C) (B) Alluvium plot visualizing the distribution of the samples according to different clusters. D) Distribution of lupus severity index across clusters with p-value computed using an ANOVA test.

We observed in CD4^+^ T cells a positive association with lupus severity index^19^, SLEDAI activity score^10,21^, ACR criteria thombocytopenia, and ACR criteria Oral Ulcers. CD4 transcriptomic cluster 1 was comprised of 44 patients and was characterized by a high prevalence of photosensitivity, seizures, younger age at diagnoses, and presence of anti-Smith autoantibody positivity (Table S4). Interestingly, we detected a positive association with CD4^+^ T cells cluster1 and lupus severity index, which is a validated tool based on weighted scores of the ACR classification criteria^19^. The ACR criteria positively associated with CD4^+^ T cells cluster1 were history of seizure and thrombocytopenia. There was also a positive association with this cluster and the SLICC score, which tracks organ damage over time, likely associated with disease severity. The same cluster was negatively associated with age at diagnoses, ACR leukopenia, ACR psychosis (Figure 2B).

CD4^+^ T transcriptomic cluster 2 was positively associated with age at diagnoses, SLEDAI score, and ACR leukopenia. The same cluster was negatively associated with ACR seizure. Patients with a high prevalence of malar rash, psychosis, anti-dsDNA autoantibody positivity, and SLEDAI score constitute the CD4 transcriptomic cluster 2 (Table S4). Finally, CD4+ T transcriptomic cluster 3 was negatively associated with ACR malar rash and presence of anti-Smith autoantibody positivity.

In CD14^+^ monocytes we observe a significant positive association in cluster 1 with White as well as a negative association with Asian (Figure 2B). Moreover, cluster 2 of the CD14^+^ monocytes was negatively associated with race White. This was the only association observed for cluster 2. On the clinical level, cluster 1 of CD14^+^ monocytes consisted of 29 patients and it was defined by lower rates of fare, and anti-dsDNA autoantibody positivity, while cluster 2 encloses 58 patients characterized by high rates of flare severity (measured as mild 1, moderate 2, severe 3) (Table S5).

The two clusters of CD19, B cells, showed for cluster 1 a negative association for SLICC score, lupus severity index, and ACR seizure, while on the other hand for cluster 2 we observe a positive association for SLICC score, lupus severity index, and ACR seizure. (Figure 2B). Cluster 1 of B cells comprised of 68 patients with a low prevalence of seizure, lymphopenia and by low score for lupus severity index and SLICC score, which suggested low disease activity. On the other hand, cluster 2 of B cells (n = 38) featured patients with high prevalence of seizure, lymphopenia and by a high score for lupus severity index and SLICC score, suggesting high disease activity (Table S6).

In the two clusters of the NK cells we observed a significant association between the presence of antinuclear antibody and cluster 1 as well as a negative association between antinuclear antibody and cluster2. No further associations were seen between the clinical and demographic parameters in the two clusters of the NK cells (Figure 2B, Table S7).

Interestingly, from this correlation analysis, we noted a similar relationship for the transcriptomic clusters of CD4+ T cells and the clinical K-means clusters previously identified by our group solely using ACR clinical criteria^12^ (Figure 2C). Supporting the similarity between clusters identified in the CD4^+^ T cells and the clustering using uniquely clinical features we identified an important overlap in patient membership (Figure 2C). A total of 61% of the individuals in the CD4^+^ T cell transcriptomic cluster 1 were also present in the severe cluster of the clinical data. Finally, as also observed in the clinical clusters, we statistically confirmed association with CD4^+^ T cells transcriptomic clusters and lupus severity index (p < 0.05) (Figure 2D). Overlaying this information allowed us to look at a global relationship between clinical data and the transcriptional profiles across the cell types. In addition, as we detected an association between CD14^+^ monocytes and ethnicity, we used DESeq2 to identify the differentially expressed genes between SLE patients from different ethnic backgrounds.

### Differences in gene expression levels within each cell type significantly differs between ethnic populations

Differential expression analysis on demographic and clinical features identified a few numbers of significant (FDR<0.05 and log2FC 1) differentially expressed genes. Specifically, 29 genes were identified for the DE on lupus severity index, 3 on SLICC score, 5 for DE on SLEDAI score, 9 for DE on ACR criteria for lupus nephritis (Table S8). Pathway analyses derived from differential expression analysis on the three different clinical indexes and the presence of lupus nephritis, a type of kidney complication caused by SLE, across the immune cell types showed enrichment of pathways in CD4 and CD14 for lupus severity index and SLICC score while SLEDAI score is enriched in CD4 only (Figure S6).

Remarkably, a larger number of differentially expressed genes between Asian and White groups were identified for each cell-type (Figure 3A, Figure S2-5). In the CD4^+^ T cells for the White cohort the Actin Filament Associated Protein 1 (*AFAP1*), *USP32P1*, and, *NAP1L3* (Nucleosome Assembly Protein 1 Like 3) were the top three significantly upregulated (padj < 0.05) genes. *ARHGEF10*, and FMN*1* were the top significant genes in CD4^+^ T cells upregulated in Asians. An example of significantly up-regulated genes in CD14^+^ monocytes cells for the White cohort were *SNORD3B-2* and the lncRNA maternally expression gene 3, *MEG3*. In Asians, Neurotensin Receptor 1 (*NTSR1*), and *RETN* were among the top significantly up-regulated genes in CD14^+^ monocytes. Top up-regulated genes in B cells for the White cohort were *ARHGAP24* and *CHL1* (Cell Adhesion Molecule L1 Like). Genes encoding for immunoglobulin heavy- and light-chain genes were statistically significantly up-regulated in Asians in B cells. Regarding NK cells, examples of upregulated significant genes in Whites were *SNORD3B-2*, and Cytokine Like 1 (*CYTL1*), while an example of up-regulated genes in Asians were *HPGD*, and *ADAM19*. Pathway analyses in Asians revealed up-regulated pathways involved in metabolism and transcriptional activity in CD14^+^ monocytes and CD4^+^T cells whereas in Whites, both B cell and NK cells were enriched for up-regulated pathways for integrin and IL2 signaling indicating a higher activity of specific cell types in Asians and Whites respectively (Figure 3B). There were no genes shared across all four cell types, (Table S2) and only *RPL3P2* is shared across CD19, CD4, and NK cells.

**Figure 3:**
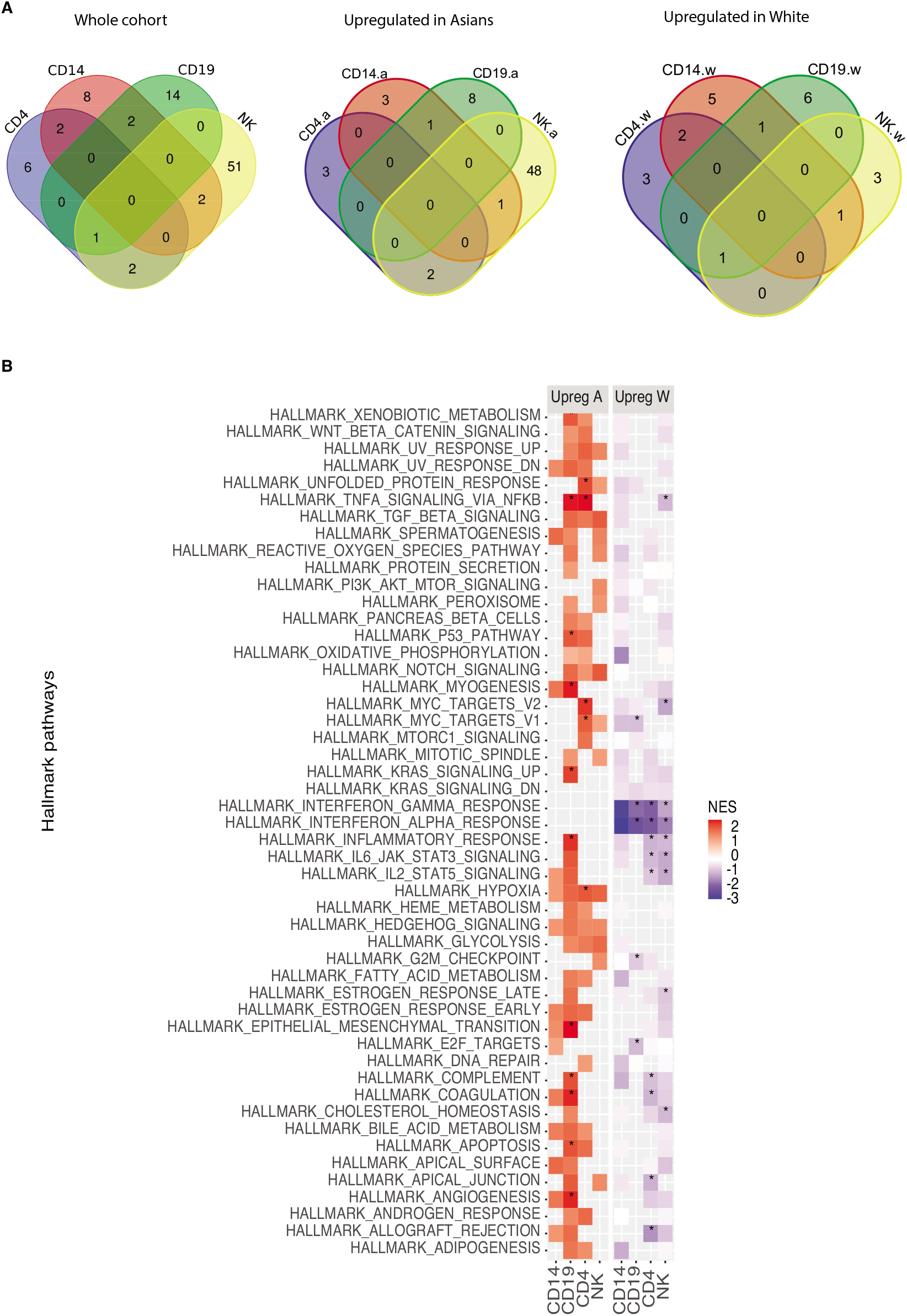
A) Identification of the common DE genes (padj < 0.05 & abs log2FC 1) White vs Asians across the different cell type. Table S2 lists the genes shared and specific for each cell type. B) Pathway enrichment analyses. for the predicted target genes of White vs Asians.

### Gene co-expression analyses within ethnic populations and disease activity revealed specific modules of genes

Classical approaches to analyze the transcriptome data by using differential gene expression analysis based on sample groups defined by a selection of clinical parameters precluded dissection of the heterogeneity of SLE. Co-expression analysis, on the other hand, identifies similarly regulated genes across samples, and then groups these genes into modules, which can then be explored for each patient sample individually or for entire patient groups^22,23^. As we found the largest number for DE genes on ethnicity, we decided to use CoCena2 on the patients’ data grouped by ethnic background combined with disease activity (SLEDAI score). Hence, patients were divided into White-high, Asian-high (high disease activity defined by a SLEDAI score >=6) from White-low, Asian-low (low disease activity defined by a SLEDAI score < 6) for each cell type. The CoCena pipeline was run on each cell line independently. The pipeline identified 8, 5, 10, and 4 co-expression modules for CD4^+^ T cells, CD14^+^ monocytes, CD19 and NK cells respectively. Analysis of the modules revealed group-specific enrichment of co-expressed gene modules (Figure 4A). Enrichment analyses on each of the modules identified associated gene signatures displaying distinct functional characteristics, which distinguish the different sample groups. (Figure 4B, Figure 5). We then annotated the modules according to their behavior into seven categories: ethnicity specific independent of disease activity, opposite behaviors according to ethnicity, White specific, Asian specific, low disease activity specific, high disease activity specific, and high disease activity specific for White. For example, the IL6 JAK STAT3 signaling pathway was identified as having opposite behavior in Asian and Caucasian populations and was enriched in the modules dark grey and gold for CD4^+^ T cell. More precisely the dark grey module was highly expressed in Asians patients with low disease activity and in White patients with high disease activity. The maroon module of CD14^+^ monocytes was annotated as being specific for low disease activity as it was low expressed in Asians and White individuals with high SLEDAI score. Interestingly, seven of the 10 modules in B cells were highly expressed in Whites compared to Asians regardless of their disease activity, while three of the four modules in the NK cells were exclusively highly expressed in Asians patients with high disease activity (Figure 4A). An extended analysis of the modules revealed a prominent enrichment of inflammatory, autoimmune, and interferon pathways (Figure 5, Figures S7).

**Figure 4:**
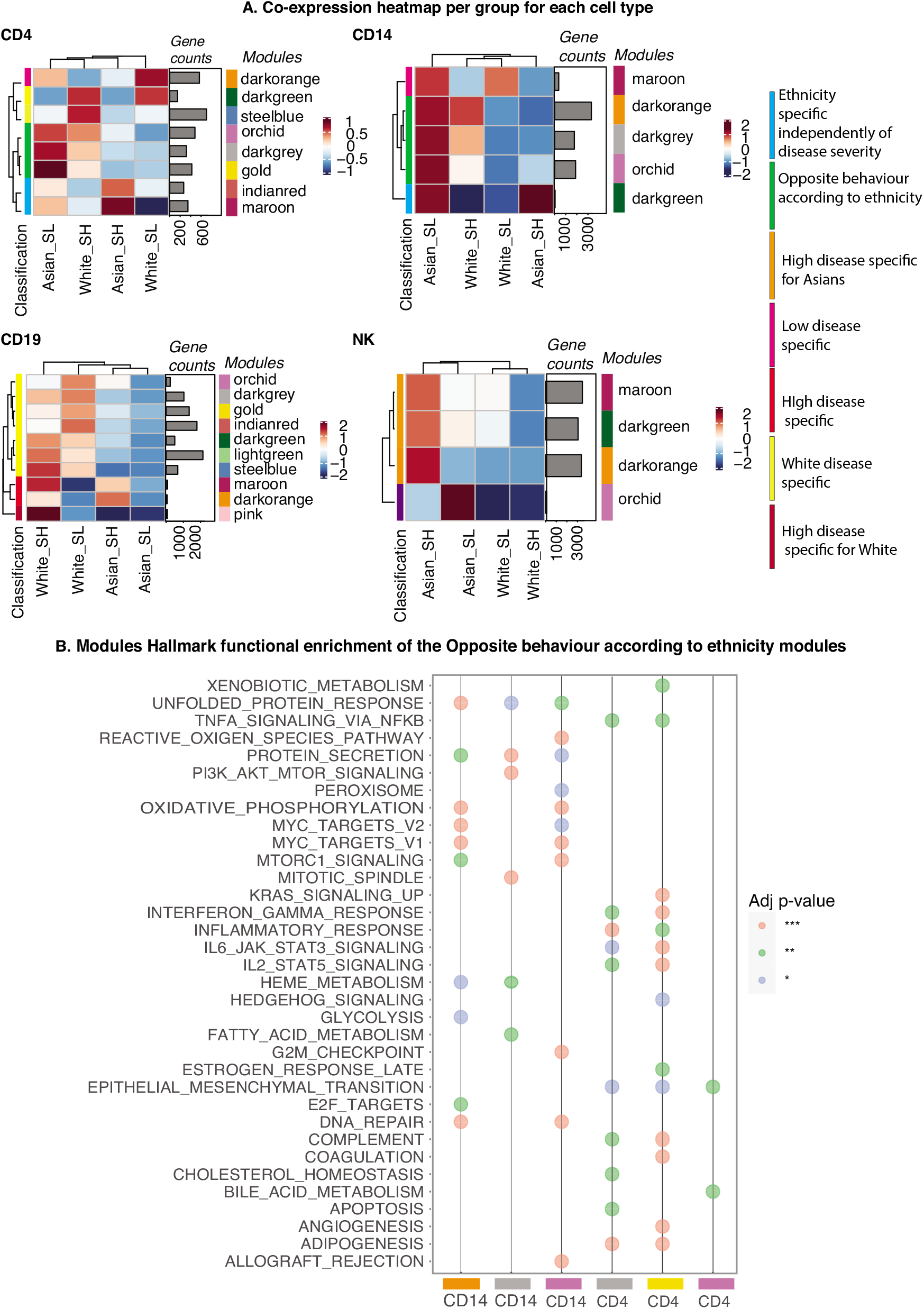
A) Group fold change heat map and hierarchical clustering for the four data-driven sample groups and the gene modules identified byCoCena2 analysis. Classification: the annotation of the modules into different categories based on their behavior. B) Functional enrichment of CoCena2-derived modules classified as “Opposite behavior according to ethnicity using the Hallmark gene set database. Only significant terms are visualized.

**Figure 5:**
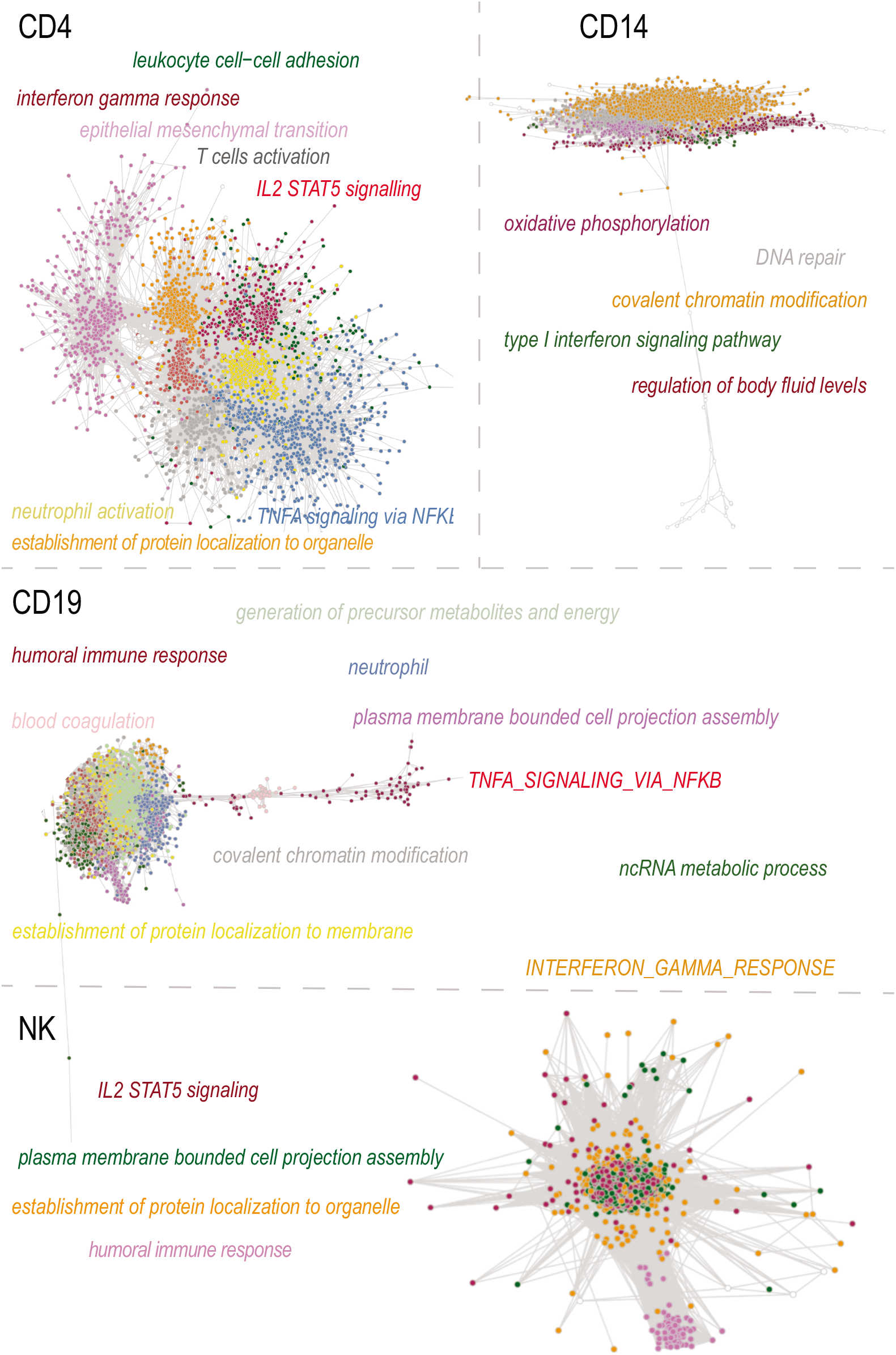
Visualization of the CD4, CD14, CD19, and NK cells CoCena2 network. Nodes are genes and edges represent co-expressed genes. Additional module information is displayed by module-colored labels. Labels include information about top-GO term as well as representative Hallmark terms.

### Machine Learning approaches predicted SLE disease activity in individual ethnic groups

Based on the results of the previous analyses and on the clustering of the patients according to co-expressed modules found on ethnicity and disease activity, we decided to use random forest (RF) on the transcriptomic data for each cell type within each racial group to discriminate between patients with high disease activity (SLEDAI score >=6) and low disease activity (SLEDAI score < 6). Receiver operating characteristic (ROC) analysis of each RF model showed an AUC of 0.8 (Figure 6A), indicating the consistent efficiency of the model in discriminating high disease activity samples from low disease activity samples in CD4^+^ T cells and CD19 cells when the whole cohort is taken into account. With regards to the White cohort, the RF model trained on CD4^+^ T cells and CD14^+^ monocytes showed an AUC of 0.71 and 0.79 respectively. On the other hand, the models trained on the Asian cohort showed an AUC higher than random only in CD4 T cells (AUC 0.62). The models trained with the top feature of the other ethnic background showed an AUC less than 0.6 which indicates inefficiency in the model in discriminating patients based on disease activity and underlaying the ethnicity specific features for each model (Figure S8).

**Figure 6:**
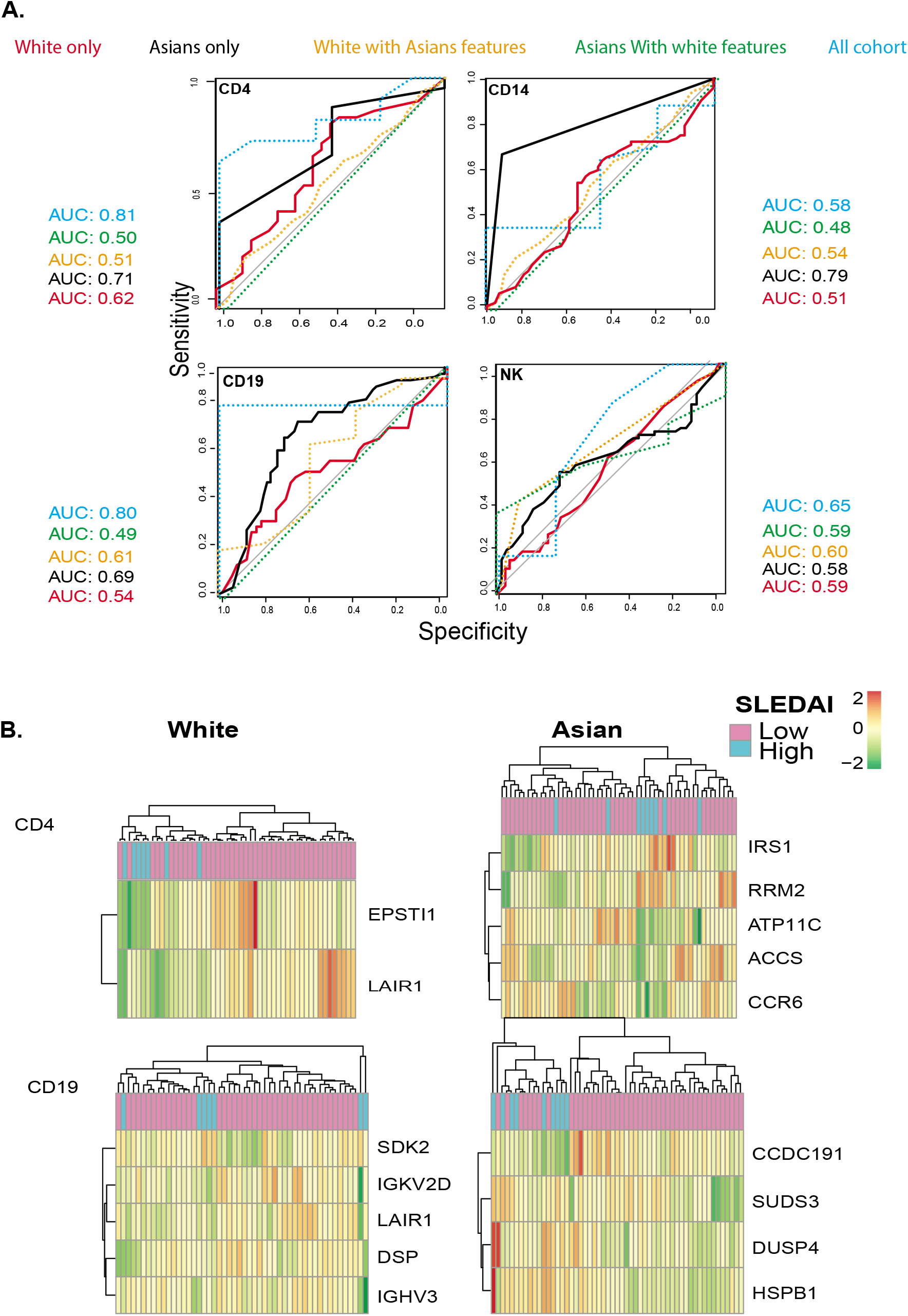
A) Area under the ROC curve of machine learning classifier across the White and Asians data sets in discriminating active vs inactive patients. White (black) and Asians (red) B) Clustergram generated by using the expression values of the top predictors selected by FSelector function.

The top contributing feature identified by the random forest algorithm to segregate high and low disease activity patients in CD4^+^ T cell in the White cohort were *EPSTI1* and *LAIR1*, while in the Asian cohort these were: *IRS1, RRM2, ATP11C, ACCS*, and *CCR6*. With regards to the B cells, the top features for the White cohort were *SDK2, IGKV2D, LAIR1, DSP*, and *IGHV3* while top features for the Asians cohort were: *CCFC292, SUDS3, DUSP4*, and *HSPB1* (Figure 6B). Importantly some of these genes have been previously associated with SLE. We observed that the model was able to select SLE relevant features. In summary, these results suggested the potential interaction between clinical features and potentially disease related genes, which could help explain the multifactorial, heterogeneous and systemic nature of the disease within White and Asians.

## Discussion

SLE has great diversity of presentation and treatment response. There has been considerable progress in genetic studies of SLE, thanks partly to technological advances and to confirming previously reported genes. Due to the complexity of the pathogenesis and the heterogeneous nature of SLE clinical manifestations, imprecise diagnosis and poor treatment efficacy remain two major obstacles that drastically affect the outcome of patients with SLE. Here we profiled a multiethnic cohort of 120 patients with SLE using cell sorting RNA-seq data (CD4^+^ T cells, CD14^+^ monocytes, B cells, and NK cells). Since SLE severity is known to vary widely between racial and ethnic groups, analysis of a large multiethnic cohort is crucial for understanding the genetic and non-genetic determinants of this ethnic-associated variability.

With an unsupervised clustering approach, we were able to unbiasedly stratify patients based on gene expression signatures allowing the identification of separate molecular pathways underpinning disease in SLE for each cell type (Figure 2A). Transcriptomic cluster 1 of the CD4^+^ T cells has a lower age of onset (mean 24 y/o) compared to cluster 2 (31 y/o) and cluster 3 (29 y/o) of the same cell type (Table S4) suggesting an enrichment of higher disease severity in cluster 1 compared to cluster 2 in CD4^+^ T cells. This is also supported by the alluvium plot showing a higher clustering of severe patients, according to k-means clustering on only clinical criteria, in the cluster 1 and 2 of CD4^+^ T cells (Figure 2C). As shown also on the clinical data work, the Lupus Severity Index, a validated scoring system based on the ACR classification criteria, was also significantly different between the three clusters (Figure 2D).

Within each immune cell-type, each identified cluster was characterized by specific clinical features (Table S4-7). We also observe an opposite association between cluster 1 and cluster 2 of the CD14^+^ monocytes and ethnicity (Asians and White). To better understand this association, we decided to conduct differential expression analysis to extract the cell-type specific transcriptional signature expression in SLE patients of ethnically different background. Table S2 lists the significant differentially expressed genes for each cell line. Some of the shared genes across the four cells were involved in SLE or autoimmune related pathways such as: *AFAP1*, which was studied in Plasmacytoid dendritic cells in lupus mice.^24^, *USP32P1* which is an Immune-Induced gene specific of Caucasians only^25^, and *RPL3P2* which is a gene mapping in the HLA class I region and in the autosomal dominant polycystic kidney disease gene region^26^. The top three up-regulated genes in the CD4^+^ T cells for the White cohort were *AFAP1*, a gene shown to have an expression pattern similar to the interferon alpha in SLE mice^24^, *USP32P1*, a ubiquitinase specific of the White population^25^, and *NAP1L3*, Nucleosome Assembly Protein 1 Like 3, which has been recently listed as one of the 93-SLE gene signature (SLE MetaSignature) that is differentially expressed in the blood of patients with SLE compared with healthy volunteers^27^. *ARHGEF10, FMN1*, and *CD79A* are the top three significant up-regulated genes in CD4^+^ T cells in Asians. These three genes are either specific for the immune system and have been shown to be associated with SLE^28,29^. Two of the top significant up-regulated genes in CD14^+^ monocytes for the White cohort were *SNORD3B-2*, a gene involved in the biological pathways regulated by progranulin^30^, and the lncRNA maternally expression gene 3, *MEG3*. Long non-coding RNAs (lncRNAs) have emerged as important regulators of biological processes and substantial evidences have been accumulated showing that lncRNAs involved in the pathogenesis of the rheumatoid diseases such as SLE^31^. In Asians, Neurotensin Receptor 1 (*NTSR1*), and *RETN*, a gene involved in the development of cardiovascular diseases among Egyptian SLE patients^32^, were between the significant up-regulated genes in Asians in CD14^+^ monocytes. Examples of up-regulated genes in B cells for the White cohort were *ARHGAP24*, shown to be upregulated in SLE patients^33^, and *CHL1* (Cell Adhesion Molecule L1 Like), a protein encoded by this gene is a member of the L1 gene family of neural cell adhesion molecules. Genes encoding for immunoglobulin heavy- and light-chain genes were statistically significant up-regulated in Asians in B cells. These genes play an important role in antigen recognition^34^. Regarding NK cells, upregulated significant genes in White were *SNORD3B-2*, and Cytokine Like 1 (*CYTL1*), while in Asians were *HPGD*, and *ADAM19*, a gene highly expressed when SLE murine models are subminiaturized with VGX-1027^35^. Gene set enrichment analyses (GSEA) on differentially expressed genes on SLICC score, SLEDAI score, lupus severity index, and lupus nephritis (Figure S6) showed and elevated expression of interferon pathways in the CD14^+^ monocytes. We observe and enrichment of interferon pathways in the CD4^+^ T cells on the differentially expressed analysis conducted on SLEDAI score.

Analysis of gene expression data has been widely used in transcriptomic studies to understand functions of molecules inside a cell and interactions among molecules, however differential gene expression analysis might prevent the dissection of the heterogeneity of the disease. For this reason, we conducted co-expression analyses using CoCena taking demographic and clinical features into account. Co-expression analysis identified similarly regulated genes across samples and grouped these genes into modules. Samples were grouped into White-high, Asians-high (high disease activity defined by a SLEDAI score >=6) and White-low, Asians-low (low disease activity defined by a SLEDAI score < 6). Modules were annotated according to their behaviors into seven categories (Figure 4A). For example, Interferon gamma response pathway was enriched in modules dark grey and gold for CD4^+^ T cell (Figure 4B). Extended analysis of the modules revealed a prominent enrichment of inflammatory, autoimmune, and interferon pathways (Figure 4 and Figure S7). Cocena2 further identified modules which contained transcription factor signature genes known to be associated with SLE. As an example the top three transcription factor (TF) in the dark orange module of CD4^+^ T cells are *ETS1*^36^, *SP1*^37^, and *ELK4*^38^, and for the gold module in the CD4^+^ T cells the top three TF are *Sfpi1-1*^39^, *SPI1*^37^, and *Hand1*^40^. We also observe interferon enriched modules in dark grey, gold, Indian red, and maroon (CD4), dark green (CD14), dark orange (CD19 and NK) (Figure S7).

Finally, using a RF approach we were able to extract features within each cell type for each ethnic group that help to discriminate high disease from low disease activity patients. Random forest classification of disease activity in the White and Asian cohorts showed the best classification in CD4^+^T cells White cohort with two genes as features (*EPSTI1* and *LAIR1*), which corresponded to an AUC value of 0.71 (Figure 6). Both genes are involved in the interferon pathways. Epithelial stromal interaction 1 (*EPSTI1*) is an interferon (IFN) response gene^41^, *LAIR-1* may play a relevant role in the mechanisms controlling IFNα production by dendritic cells both in normal and pathological innate immune responses^42^, *IRS1* expression is altered in SLE mice^43^, and the frequency of the *IGHV3* gene family has been assessed over glucocorticoid time treatment^44^. This was the strongest results observed suggesting that genes expressed by CD4^+^T cells may prove to be informative in the study of cell-specific methods of SLE pathogenesis.

Strengths of this study include the rich phenotyping data and one of the largest SLE cohorts including White, and Asian patients to be profiled for cell sorted RNA-seq, which allowed us to identify genetic effects of race in SLE disease activity, shedding light on molecular mediators of race in disease heterogeneity. Future work will include testing these findings in other multi-ethnic cohorts and include a multi-omics approach. There are several limitations that should be recognized. Our study is limited by the restriction of transcriptomic data to one time point. At the time of blood collection, patients were not currently in lupus flares. However, since gene expression is dynamic and may reflect cellular responses to an underlying disease process, it is possible that patients transition between transcriptomic clusters as during their disease course. This remains a testable hypothesis in future studies incorporating longitudinal profiling. Additionally, specificity and sensitivity were low in all cell types and this could be due to the imbalanced dataset. ML models with substantial predictive accuracy can assist clinicians with complicated diagnostic decision making, though extensive study is necessary to construct accurate models for specific population groups. Our study is also limited by the fact that all patients within our cohort are currently stable on medication making the identification of transcriptomic differences between patients nontrivial as well as the absence of controls. Although there are large numbers of publicly available gene expression profiles of SLE patients, many of these profiles are not annotated with SLEDAI data and are not cell sorted data and do not examine specific cell types. Furthermore, some data sets which include SLEDAI data show heavy class imbalance, which impedes classification. Further work to integrate cross-platform expression data will be crucial to expanding our ability to classify active and inactive SLE patients.

In summary, we have identified distinct clinical subtypes of SLE using immune cell sorted transcriptomic data that have distinct associations with clinical measures, specifically with lupus severity index. We observe a relationship between transcriptomic clusters of CD4^+^ T cells and the clinical K-means clusters of the same patients. We also identified a positive association between the transcriptomic cluster 1 of the CD14^+^ monocyte cells and White ethnicity. Pathway enrichment analyses on the differential expressed genes highlighted cell type specific upregulated immune pathways for either Asian or White SLE patients including TNF-alpha signaling via NFKB in CD4^+^T cells in Whites and IL2 STAT5 signaling in NK cells in Asians. Additionally, we are able to identify modules of genes that are either ethnicity specific, disease specific as well as train machine learning models to discriminate patients based on disease activity within each ethnic group. Our findings provide an insight of the cellular processes that drive SLE pathogenesis within patients of different ethnicity and may eventually lead to customized therapeutic strategies based on patients’ unique patterns of cellular activation.

## Methods

### Study Design

Participants were recruited from the California Lupus Epidemiology Study (CLUES). CLUES was approved by the Institutional Review Board of the University of California, San Francisco. All participants signed a written informed consent to participate in the study. Study procedures involved an in-person research clinic visit, which included collection and review of medical records prior to the visit; a history and physical examination conducted by a physician specializing in lupus; collection of biospecimens, including peripheral blood for clinical and research purposes; and completion of a structured interview administered by an experienced research assistant. All SLE diagnoses were confirmed by study physicians based upon one of the following definitions: (a) meeting□≥□4 of the 11 American College of Rheumatology (ACR) revised criteria for the classification of SLE as defined in 1982 and updated in 1997, (b) meeting 3 of the 11 ACR criteria plus a documented rheumatologist’s diagnosis of SLE, or (c) a confirmed diagnosis of lupus nephritis, defined as fulfilling the ACR renal classification criterion (>0.5 grams of proteinuria per day or 3□+□protein on urine dipstick analysis) or having evidence of lupus nephritis on kidney biopsy. A total of 120 patients (Table S1) were profiled with bulk RNA-seq from the CLUES cohort.

### RNA-seq processing and quality control

Peripheral blood mononuclear cells were isolated from patient donors. Cells were isolated from peripheral blood utilizing magnetic beads (CD14^+^monocytes, B cells, CD4^+^T cells and NK cells) using EasySep protocol from STEM cell technologies on 120 patients. RNA and DNA were extracted using Qiagen column and quality was assessed on Agilent bioanalyzer. 1ng - 5ng of RNA was used with SmartSeqv2 protocol to get cDNA, followed by Illumina Nextera XT DNA library prep with input of 0.8ng cDNA and 15 cycles of PCR amplification. Both cDNA and DNA libraries qualities were controlled on Agilent bioanalyzer. Libraries were sequenced on HiSeq4000 PE150.

Salmon v0.8.2 was used for our alignment-free pipeline. Adaptor-trimmed reads were used as input. We quantified gene expression using raw counts and kept the genes that showed an average FPKM value across the samples >0.5. FastQC (v0.11.8) was run to assess the quality of the sequence reads. To further assess for low quality samples we checked the expression of 10 housekeeping genes (e.g. *SNRPD3, EMC7*, and *VPS29*) called the Eisenberg housekeeping genes^16^. Finally, we conducted K-means clustering and removed all outlier samples that were > 1 standard deviation from the center of the cluster. The Davies-Bouldin’s index was also computed to confirm cluster stability. Batch correction was performed with limma (v 3.40.6).

### Unsupervised clustering approach and association analyses

Unsupervised K-means clustering was performed on the batch corrected transcriptomic data per cell type using the factoextra package (v 1.0.7). The number of clusters, k, was chosen by maximizing cluster stability measured by Jaccard similarity using a bootstrap resampling-based method. The identified clusters were then tested for associations with clinical variables such as sex, race, and the ACR criteria. Fisher’s exact test was used to evaluate if categorical variables were enriched in a cluster with respect to the others. For continuous variables, we used ANOVA testing.

### Differential expression analyses

We performed differential expression gene testing with DESeq2 (v.1.24.0 R package) using default settings. Sequence lane, medications, and sex were used as covariates within the DESeq2 model. Statistical significance was set a 5% false discovery rate (FDR; Benjamini-Hochberg). The bioconductor package fgsea (v 1.10.1), was used for gene set enrichment analysis (GSEA). Differential expression analysis was conducted for several clinical features, specifically: SLEDAI score^17^, the mean Systemic Lupus International Collaborating Clinics/American College of Rheumatology (SLICC) Damage Index score^18^, lupus severity index, a validated scoring system based on the ACR classification criteria^19^, the presence of lupus nephritis, and ethnicity (White vs Asians), Figure S2-6.

### Co-expression analysis

The CoCena2 pipeline [https://github.com/Ulas-lab/CoCena2] was used to define differences and similarities in transcript expression patterns between the two ethnic groups and disease activity (based on the SLEDAI score). Patients were divided in White-high, Asians-high (high disease activity defined by a SLEDAI score >=6) from White-low, Asians-low (low disease activity defined by a SLEDAI score < 6) for each cell type. As previously described in other publications, CoCena2 (Construction of co-expression Network Analysis - automated) was performed based on Pearson correlation. All genes were used as the input. Pearson correlation was performed using the R package Hmisc (v4.1-1). To increase data quality, only significant (p < 0.05) correlation values were kept. The nodes were colored based on the Group Fold Change (GFC), the mean of each condition versus overall mean for each gene respectively, for each group separately. Unbiased clustering was performed using the “label propagation” algorithm in graph (v1.2.1) and was repeated 1000 times. Genes assigned to more than 5 different clusters during the iterations received no cluster assignment. The mean GFC expression for each cluster and condition were visualized in the Cluster/Condition heatmap. Clusters smaller than 10 genes were not shown^20^.

### Machine learning

Prior to applying machine learning, gene selection and normalization were performed using the R packages DaMIRSEq. (v1.2.0) and MLSeq (v 2.2.1). Within each cell type, the data was split into training and testing set at 70% for training and 30% for testing in order to distinguish high disease activity (SLEDAI score >= 6) samples from low disease activity samples (SLEDAI score < 6) in both the Asian and White populations. To prevent the inclusion of redundant features that may decrease the model performance during the classification step, a function that produces a pair-wise absolute correlation matrix was applied. The FSelector package was used to rank features. For each random forest model, we calculated the mean area under the curve (AUC) over ten cross validation folds. As a cross validation, features identified in the White cohort for a specific cell type were used to train the model on the Asian cohort for the same cell type and vice versa. Mean AUC was assessed over ten cross validation folds.

## Acknowledgements

We are grateful to all of the patients who participated in the study. We are also grateful to Ms. Bushra Samad, and Dr. Ellie Seaby who provided advice on QC methods and discussion of study results.

This study was funded through the following grants: P30 AR070155 (M.S., G.A., L.A.C.), P60AR053308 (L.A.C.), K01LM012381 (M.S.), F32 AR070585 NIAMS (M.G.), U01DP005120 CDC (L.A.C., C.M.L., J.Y., M.D., L.T., P.K.), the Rheumatology Research Foundation 128849A (C.M.L.), and the Lupus Research Alliance (L.A.C.).

## Author contribution

GA: study design, data analysis, quality control analyses, data interpretation, and paper preparation.

CML, IP, TSJ, AJC, CJY: data interpretation, and paper revision.

CJY, JN, KET, LM: data generation, quality control analyses, genetic ancestry estimates, and paper revision.

PK, MD, JY: CLUES co-investigators. Patient enrollment and clinical, demographic characterization of patients and paper revisions.

LAC: data generation, study design, data interpretation and paper revisions.

MS: Computational expertise, study design, data interpretation and paper revisions.

