## Supplementary material for "Ethnicity-specific transcriptomic variation in immune cells and correlation with disease activity in systemic lupus erythematosus": supp figures and tables

**Supplementary Tables:**

|  |  | **Asian** | **White** | **Total** | **p value*** |
| --- | --- | --- | --- | --- | --- |
| **Number of Subjects** |  | **64** | **56** | **120** |  |
| **Female** | Female | 58 (90.6%) | 48 (85.7%) | 106 (88.3%) | 0.403 |
|  | Male | 6 (9.4%) | 8 (14.3%) | 14 (11.7%) |  |
| Lupus severity Index | Mean (SD) | 7.075 (1.621) | 6.333 (1.637) | 6.729 (1.664) | 0.014 |
|  | Range | 3.825 - 9.497 | 3.578 - 8.983 | 3.578 - 9.497 |  |
| **SlLICC score** | Mean (SD) | 1.031 (1.425) | 1.125 (1.652) | 1.075 (1.529) | 0.739 |
|  | Range | 0.000 - 6.000 | 0.000 - 6.000 | 0.000 - 6.000 |  |
| SLEDAI-2K | Mean (SD) | 2.719 (2.628) | 2.589 (3.190) | 2.658 (2.892) | 0.808 |
|  | Range | 0.000 - 11.000 | 0.000 - 16.000 | 0.000 - 16.000 |  |
| Age at diagnoses | Mean (SD) | 27.531 (12.269) | 28.661 (11.752) | 28.058 (11.994) | 0.609 |
|  | Range | 9.000 - 62.000 | 9.000 - 64.000 | 9.000 - 64.000 |  |
| **ACR Malar Rash** | Mean (SD) | 0.422 (0.498) | 0.464 (0.503) | 0.442 (0.499) | 0.644 |
|  | Range | 0.000 - 1.000 | 0.000 - 1.000 | 0.000 - 1.000 |  |
|  | N positive | 27 | 26 | 53 |  |
| ACR Discoid Rash | Mean (SD) | 0.109 (0.315) | 0.107 (0.312) | 0.108 (0.312) | 0.969 |
|  | Range | 0.000 - 1.000 | 0.000 - 1.000 | 0.000 - 1.000 |  |
|  | N positive | 6 | 7 | 13 |  |
| **ACR Oral Ulcers** | Mean (SD) | 0.469 (0.503) | 0.661 (0.478) | 0.558 (0.499) | 0.035 |
|  | Range | 0.000 - 1.000 | 0.000 - 1.000 | 0.000 - 1.000 |  |
|  | N | 29 | 38 | 67 |  |
| ACR Photosensitivity | Mean (SD) | 0.312 (0.467) | 0.446 (0.502) | 0.375 (0.486) | 0.133 |
|  | Range | 0.000 - 1.000 | 0.000 - 1.000 | 0.000 - 1.000 |  |
|  | N positive |  | 27 |  |  |
| **ACR Arthritis** | Mean (SD) | 0.703 (0.460) | 0.857 (0.353) | 0.775 (0.419) | 0.044 |
|  | Range | 0.000 - 1.000 | 0.000 - 1.000 | 0.000 - 1.000 |  |
|  | N positive | 45 | 48 | 93 |  |
| ACR Serositis | Mean (SD) | 0.312 (0.467) | 0.518 (0.504) | 0.408 (0.494) | 0.022 |
|  | Range | 0.000 - 1.000 | 0.000 - 1.000 | 0.000 - 1.000 |  |
|  | N positive | 19 | 29 | 48 |  |
| **ACR Pleuritis** | Mean (SD) | 0.219 (0.417) | 0.446 (0.502) | 0.325 (0.470) | 0.008 |
|  | Range | 0.000 - 1.000 | 0.000 - 1.000 | 0.000 - 1.000 |  |
|  | N positive | 14 | 25 | 39 |  |
| ACR Pericarditis | Mean (SD) | 0.156 (0.366) | 0.143 (0.353) | 0.150 (0.359) | 0.839 |
|  | Range | 0.000 - 1.000 | 0.000 - 1.000 | 0.000 - 1.000 |  |
|  | N positive | 10 | 8 | 18 |  |
| **ACR Neurologic** | Mean (SD) | 0.062 (0.244) | 0.143 (0.353) | 0.100 (0.301) | 0.146 |
|  | Range | 0.000 - 1.000 | 0.000 - 1.000 | 0.000 - 1.000 |  |
| ACR Seizure | Mean (SD) | 0.016 (0.125) | 0.107 (0.312) | 0.058 (0.235) | 0.033 |
|  | Range | 0.000 - 1.000 | 0.000 - 1.000 | 0.000 - 1.000 |  |
|  | N positive | 2 | 5 | 7 |  |
| **ACR Psychosis** | Mean (SD) | 0.047 (0.213) | 0.036 (0.187) | 0.042 (0.201) | 0.763 |
|  | Range | 0.000 - 1.000 | 0.000 - 1.000 | 0.000 - 1.000 |  |
|  | N positive | 3 | 2 | 5 |  |
| ACR Immunologic | Mean (SD) | 0.857 (0.363) | 0.736 (0.443) | 0.750 (0.435) | 0.208 |
|  | Range | 0.000 - 1.000 | 0.000 - 1.000 | 0.000 - 1.000 |  |
| **ACR anti-dsDNA** | Mean (SD) | 0.719 (0.453) | 0.464 (0.503) | 0.600 (0.492) | 0.004 |
|  | Range | 0.000 - 1.000 | 0.000 - 1.000 | 0.000 - 1.000 |  |
|  | N positive | 47 | 25 | 72 |  |
| ACR anti-Smith | Mean (SD) | 0.297 (0.460) | 0.214 (0.414) | 0.258 (0.440) | 0.306 |
|  | Range | 0.000 - 1.000 | 0.000 - 1.000 | 0.000 - 1.000 |  |
|  | N positive | 19 | 12 | 31 |  |
| **ACR ANA** | Mean (SD) | 0.969 (0.175) | 0.946 (0.227) | 0.958 (0.201) | 0.545 |
|  | Range | 0.000 - 1.000 | 0.000 - 1.000 | 0.000 - 1.000 |  |
|  | N positive | 61 | 54 | 115 |  |
| ACR Hematologic | Mean (SD) | 0.453 (0.502) | 0.411 (0.496) | 0.433 (0.498) | 0.643 |
|  | Range | 0.000 - 1.000 | 0.000 - 1.000 | 0.000 - 1.000 |  |
|  | N positive | 29 | 23 |  |  |
| **ACR Hemolytic Anemia** | Mean (SD) | 0.078 (0.270) | 0.036 (0.187) | 0.058 (0.235) | 0.327 |
|  | Range | 0.000 - 1.000 | 0.000 - 1.000 | 0.000 - 1.000 |  |
|  | N positive | 5 | 2 | 7 |  |
| ACR Leukopenia | Mean (SD) | 0.188 (0.393) | 0.196 (0.401) | 0.192 (0.395) | 0.902 |
|  | Range | 0.000 - 1.000 | 0.000 - 1.000 | 0.000 - 1.000 |  |
|  | N positive | 12 | 11 | 23 |  |
| **ACR Lymphopenia** | Mean (SD) | 0.266 (0.445) | 0.286 (0.456) | 0.275 (0.448) | 0.808 |
|  | Range | 0.000 - 1.000 | 0.000 - 1.000 | 0.000 - 1.000 |  |
|  | N positive | 17 | 16 | 33 |  |
| ACR Thrombocytopenia | Mean (SD) | 0.188 (0.393) | 0.125 (0.334) | 0.158 (0.367) | 0.354 |
|  | Range | 0.000 - 1.000 | 0.000 - 1.000 | 0.000 - 1.000 |  |
|  | N positive | 12 | 7 | 19 |  |
| **ACR.Renal** | Mean (SD) | 0.562 (0.500) | 0.304 (0.464) | 0.442 (0.499) | 0.004 |
|  | Range | 0.000 - 1.000 | 0.000 - 1.000 | 0.000 - 1.000 |  |
|  | N positive | 36 | 17 | 53 |  |
| ACR.Photosensitivity | Mean (SD) | 0.312 (0.467) | 0.464 (0.503) | 0.383 (0.488) | 0.089 |
|  | Range | 0.000 - 1.000 | 0.000 - 1.000 | 0.000 - 1.000 |  |
|  | N positive | 19 | 27 | 46 |  |
| **ACR.APLA** | Mean (SD) | 0.250 (0.436) | 0.304 (0.464) | 0.275 (0.448) | 0.516 |
|  | Range | 0.000 - 1.000 | 0.000 - 1.000 | 0.000 - 1.000 |  |
|  | N positive | 16 | 17 | 33 |  |
| ACR lupus nephritis | Mean (SD) | 0.562 (0.500) | 0.286 (0.456) | 0.433 (0.498) | 0.002 |
|  | Range | 0.000 - 1.000 | 0.000 - 1.000 | 0.000 - 1.000 |  |
|  | N positive | 19 | 27 | 53 |  |
| **Flare severity**  **(mild 1, moderate 2, severe 3)** | N-Miss | 43 | 36 | 79 |  |
|  | Mean (SD) | 1.762 (0.700) | 1.900 (0.718) | 1.829 (0.704) | 0.537 |
|  | Range | 1.000 - 3.000 | 1.000 - 3.000 | 1.000 - 3.000 |  |

**Table S1:** CLUES Cohort Demographics.

**P value done with chi-squared or ANOVA were appropriate. All are ACR clinical feature are a yes/no variables, except flare, which has 3 levels (mild 1, moderate 2, severe 3). N positive= number of individuals having the clinical feature.*

a. **Whole cohort**

| Cell | Total genes | Genes |
| --- | --- | --- |
| CD19 CD4 NK | 1 | RPL3P2 |
| CD14 CD4 | 2 | USP32P1 AFAP1 |
| CD4 NK | 2 | FMN1 RP11-345P4.6 |
| CD14 CD19 | 2 | NCF1C PF4 |
| CD14 NK | 2 | LGALS3BP SNORD3B-2 |

b. **Asian cohort**

| Cell | Total genes | Genes |
| --- | --- | --- |
| CD4 NK | 2 | FMN1 RP11 |
| CD14 CD19 | 1 | NCF1C |
| CD14 NK | 1 | LGALS3BP |
| CD4 | 3 | ARHGEF10 IGLC2 CD79A |

c. **White cohort**

| Cell | Total genes | Genes |
| --- | --- | --- |
| CD19 CD4 NK | 1 | RPL3P2 |
| CD14 CD4 | 2 | AFAP1 USP32P1 |
| CD14 CD19 | 1 | PF4 |
| CD14 NK | 1 | SNORD3B-2 |

**Table S2:** Shared genes across cell types in the differential expression analysis results for White vs Asians. Only shared genes with an adjusted p value less than 0.05 and logFC 1 shown.

| **cells/clusters** | **CD4** | **CD14** | **CD19** | **NK** |
| --- | --- | --- | --- | --- |
| **2** | 0.7919779 0.5946857 | 0.6616488 0.4648584 | 0.6427347 0.9212637 | 0.9795288 0.9056403 |
| **3** | 0.9946251 0.7046017 0.7593670 | 0.6023904 0.7393057 0.9581677 | 0.5815782 0.7681514 0.4995909 | 0.5716806 0.5427028 0.8870137 |
| **4** | 0.9828838 0.4947990 0.5699967 0.7035057 | 0.7335078 0.7570003 0.4257458 1.0000000 | 0.5758475 0.5852199 0.4596246 0.7176555 | 0.6093179 0.5381911 0.4618591 0.7202778 |
| **5** | 0.9390328 0.5067186 0.5987792 0.6810699 0.6140282 | 0.8046264 0.7701437 0.5786609 0.9293843 0.6500000 | 0.6170264 0.5799071 0.4771932 0.7763333 0.2852434 | 0.6331669 0.6113674 0.5034669 0.8259372 0.7746111 |

**Table S3:** Jaccard stability index done on *100 bootstrap iterations

| CD4 | **1 (N=44)** | **2 (N=40)** | **3 (N=27)** | **Total (N=111)** |
| --- | --- | --- | --- | --- |
| **ACR.Malar.Rash** |  |  |  |  |
| Mean (SD) | 0.477 (0.505) | 0.550 (0.504) | 0.259 (0.447) | 0.450 (0.500) |
| Range | 0.000 - 1.000 | 0.000 - 1.000 | 0.000 - 1.000 | 0.000 - 1.000 |
| N positive | 21 | 22 | 7 |  |
| **ACR.Discoid.Rash** |  |  |  |  |
| Mean (SD) | 0.091 (0.291) | 0.175 (0.385) | 0.037 (0.192) | 0.108 (0.312) |
| Range | 0.000 - 1.000 | 0.000 - 1.000 | 0.000 - 1.000 | 0.000 - 1.000 |
| N positive | 4 | 7 | 1 |  |
| **ACR.Oral.Ulcers** |  |  |  |  |
| Mean (SD) | 0.477 (0.505) | 0.650 (0.483) | 0.444 (0.506) | 0.532 (0.501) |
| Range | 0.000 - 1.000 | 0.000 - 1.000 | 0.000 - 1.000 | 0.000 - 1.000 |
| N positive | 21 | 26 | 12 |  |
| **ACR.Arthritis** |  |  |  |  |
| Mean (SD) | 0.773 (0.424) | 0.750 (0.439) | 0.741 (0.447) | 0.757 (0.431) |
| Range | 0.000 - 1.000 | 0.000 - 1.000 | 0.000 - 1.000 | 0.000 - 1.000 |
| N positive | 34 | 30 | 20 |  |
| **ACR.Pleuritis** |  |  |  |  |
| Mean (SD) | 0.273 (0.451) | 0.350 (0.483) | 0.333 (0.480) | 0.315 (0.467) |
| Range | 0.000 - 1.000 | 0.000 - 1.000 | 0.000 - 1.000 | 0.000 - 1.000 |
| N positive | 12 | 14 | 9 |  |
| **ACR.Pericarditis** |  |  |  |  |
| Mean (SD) | 0.136 (0.347) | 0.225 (0.423) | 0.111 (0.320) | 0.162 (0.370) |
| Range | 0.000 - 1.000 | 0.000 - 1.000 | 0.000 - 1.000 | 0.000 - 1.000 |
| N positive | 6 | 9 | 3 |  |
| **ACR.Seizure** |  |  |  |  |
| Mean (SD) | 0.136 (0.347) | 0.000 (0.000) | 0.037 (0.192) | 0.063 (0.244) |
| Range | 0.000 - 1.000 | 0.000 - 0.000 | 0.000 - 1.000 | 0.000 - 1.000 |
| N positive | 6 | 0 | 1 |  |
| **ACR.Psychosis** |  |  |  |  |
| Mean (SD) | 0.000 (0.000) | 0.100 (0.304) | 0.037 (0.192) | 0.045 (0.208) |
| Range | 0.000 - 0.000 | 0.000 - 1.000 | 0.000 - 1.000 | 0.000 - 1.000 |
| N positive | 0 | 4 | 1 |  |
| **ACR.anti.dsDNA** |  |  |  |  |
| Mean (SD) | 0.568 (0.501) | 0.625 (0.490) | 0.556 (0.506) | 0.586 (0.495) |
| Range | 0.000 - 1.000 | 0.000 - 1.000 | 0.000 - 1.000 | 0.000 - 1.000 |
| N positive | 25 | 25 | 15 |  |
| **ACR.anti.Smith** |  |  |  |  |
| Mean (SD) | 0.318 (0.471) | 0.325 (0.474) | 0.074 (0.267) | 0.261 (0.441) |
| Range | 0.000 - 1.000 | 0.000 - 1.000 | 0.000 - 1.000 | 0.000 - 1.000 |
| N positive | 14 | 13 | 2 |  |
| **ACR.ANA** |  |  |  |  |
| Mean (SD) | 0.977 (0.151) | 0.950 (0.221) | 0.963 (0.192) | 0.964 (0.187) |
| Range | 0.000 - 1.000 | 0.000 - 1.000 | 0.000 - 1.000 | 0.000 - 1.000 |
| N positive | 43 | 38 | 26 |  |
| **ACR.Hemolytic.Anemia** |  |  |  |  |
| Mean (SD) | 0.068 (0.255) | 0.050 (0.221) | 0.074 (0.267) | 0.063 (0.244) |
| Range | 0.000 - 1.000 | 0.000 - 1.000 | 0.000 - 1.000 | 0.000 - 1.000 |
| N positive | 3 | 2 | 2 |  |
| **ACR.Leukopenia** |  |  |  |  |
| Mean (SD) | 0.068 (0.255) | 0.325 (0.474) | 0.222 (0.424) | 0.198 (0.400) |
| Range | 0.000 - 1.000 | 0.000 - 1.000 | 0.000 - 1.000 | 0.000 - 1.000 |
| N positive | 3 | 13 | 6 |  |
| **ACR.Lymphoenia** |  |  |  |  |
| Mean (SD) | 0.295 (0.462) | 0.250 (0.439) | 0.296 (0.465) | 0.279 (0.451) |
| Range | 0.000 - 1.000 | 0.000 - 1.000 | 0.000 - 1.000 | 0.000 - 1.000 |
| N positive | 13 | 10 | 8 |  |
| **ACR.Thrombocytopenia** | |  |  |  |
| Mean (SD) | 0.250 (0.438) | 0.100 (0.304) | 0.111 (0.320) | 0.162 (0.370) |
| Range | 0.000 - 1.000 | 0.000 - 1.000 | 0.000 - 1.000 | 0.000 - 1.000 |
| N positive | 11 | 4 | 3 |  |
| **ACR.Renal** |  |  |  |  |
| Mean (SD) | 0.523 (0.505) | 0.375 (0.490) | 0.407 (0.501) | 0.441 (0.499) |
| Range | 0.000 - 1.000 | 0.000 - 1.000 | 0.000 - 1.000 | 0.000 - 1.000 |
| N positive | 23 | 15 | 11 |  |
| **ACR.Photosensitivity** |  |  |  |  |
| Mean (SD) | 0.295 (0.462) | 0.450 (0.504) | 0.407 (0.501) | 0.378 (0.487) |
| Range | 0.000 - 1.000 | 0.000 - 1.000 | 0.000 - 1.000 | 0.000 - 1.000 |
| N positive | 13 | 18 | 11 |  |
| **ACR.APLA** |  |  |  |  |
| Mean (SD) | 0.273 (0.451) | 0.350 (0.483) | 0.222 (0.424) | 0.288 (0.455) |
| Range | 0.000 - 1.000 | 0.000 - 1.000 | 0.000 - 1.000 | 0.000 - 1.000 |
| N positive | 12 | 14 | 6 |  |
| **lupus severity index** |  |  |  |  |
| Mean (SD) | 7.119 (1.567) | 6.490 (1.761) | 6.479 (1.662) | 6.736 (1.676) |
| Range | 3.825 - 9.365 | 3.825 - 8.983 | 3.578 - 9.497 | 3.578 - 9.497 |
| **slicc score** |  |  |  |  |
| Mean (SD) | 1.523 (1.849) | 0.850 (1.231) | 0.926 (1.412) | 1.135 (1.564) |
| Range | 0.000 - 6.000 | 0.000 - 5.000 | 0.000 - 5.000 | 0.000 - 6.000 |
| **sledai score** |  |  |  |  |
| Mean (SD) | 2.205 (2.511) | 3.325 (3.238) | 2.111 (2.913) | 2.586 (2.915) |
| Range | 0.000 - 9.000 | 0.000 - 16.000 | 0.000 - 11.000 | 0.000 - 16.000 |
| **acr lupus nephritis** |  |  |  |  |
| Mean (SD) | 0.500 (0.506) | 0.375 (0.490) | 0.407 (0.501) | 0.432 (0.498) |
| Range | 0.000 - 1.000 | 0.000 - 1.000 | 0.000 - 1.000 | 0.000 - 1.000 |
| **flare severity** |  |  |  |  |
| N-Miss | 29 | 25 | 20 | 74 |
| Mean (SD) | 2.000 (0.756) | 1.800 (0.676) | 2.000 (0.577) | 1.919 (0.682) |
| Range | 1.000 - 3.000 | 1.000 - 3.000 | 1.000 - 3.000 | 1.000 - 3.000 |
| **age at diagnoses** |  |  |  |  |
| Mean (SD) | 24.568 (12.043) | 30.950 (12.331) | 29.778 (11.144) | 28.135 (12.190) |
| Range | 9.000 - 59.000 | 9.000 - 64.000 | 13.000 - 50.000 | 9.000 - 64.000 |

**Table S4: clinical and demographic variables across CD4 clusters.** Flare severity measurement: mild 1, moderate 2, severe 3. All are ACR clinical feature are a yes/no variables, except flare, which has 3 levels (mild 1, moderate 2, severe 3). N positive= number of individuals having the clinical feature.

| **CD14** | **1 (N=29)** | **2 (N=58)** | **3 (N=21)** | **Total (N=108)** |
| --- | --- | --- | --- | --- |
| **ACR.Malar.Rash** |  |  |  |  |
| Mean (SD) | 0.483 (0.509) | 0.431 (0.500) | 0.429 (0.507) | 0.444 (0.499) |
| Range | 0.000 - 1.000 | 0.000 - 1.000 | 0.000 - 1.000 | 0.000 - 1.000 |
| N positive | 14 | 25 | 9 |  |
| **ACR.Discoid.Rash** |  |  |  |  |
| Mean (SD) | 0.172 (0.384) | 0.069 (0.256) | 0.143 (0.359) | 0.111 (0.316) |
| Range | 0.000 - 1.000 | 0.000 - 1.000 | 0.000 - 1.000 | 0.000 - 1.000 |
| N positive | 5 | 4 | 3 |  |
| **ACR.Oral.Ulcers** |  |  |  |  |
| Mean (SD) | 0.552 (0.506) | 0.466 (0.503) | 0.714 (0.463) | 0.537 (0.501) |
| Range | 0.000 - 1.000 | 0.000 - 1.000 | 0.000 - 1.000 | 0.000 - 1.000 |
| N positive | 16 | 27 | 15 |  |
| **ACR.Arthritis** |  |  |  |  |
| Mean (SD) | 0.828 (0.384) | 0.793 (0.409) | 0.762 (0.436) | 0.796 (0.405) |
| Range | 0.000 - 1.000 | 0.000 - 1.000 | 0.000 - 1.000 | 0.000 - 1.000 |
| N positive | 24 | 46 | 16 |  |
| **ACR.Pleuritis** |  |  |  |  |
| Mean (SD) | 0.241 (0.435) | 0.345 (0.479) | 0.381 (0.498) | 0.324 (0.470) |
| Range | 0.000 - 1.000 | 0.000 - 1.000 | 0.000 - 1.000 | 0.000 - 1.000 |
| N positive | 7 | 20 | 8 |  |
| **ACR.Pericarditis** |  |  |  |  |
| Mean (SD) | 0.172 (0.384) | 0.138 (0.348) | 0.095 (0.301) | 0.139 (0.347) |
| Range | 0.000 - 1.000 | 0.000 - 1.000 | 0.000 - 1.000 | 0.000 - 1.000 |
| N positive | 5 | 8 | 2 |  |
| **ACR.Seizure** |  |  |  |  |
| Mean (SD) | 0.103 (0.310) | 0.052 (0.223) | 0.000 (0.000) | 0.056 (0.230) |
| Range | 0.000 - 1.000 | 0.000 - 1.000 | 0.000 - 0.000 | 0.000 - 1.000 |
| N positive | 3 | 3 | 0 |  |
| **ACR.Psychosis** |  |  |  |  |
| Mean (SD) | 0.000 (0.000) | 0.052 (0.223) | 0.048 (0.218) | 0.037 (0.190) |
| Range | 0.000 - 0.000 | 0.000 - 1.000 | 0.000 - 1.000 | 0.000 - 1.000 |
| N positive | 0 | 3 | 1 |  |
| **ACR.anti.dsDNA** |  |  |  |  |
| Mean (SD) | 0.552 (0.506) | 0.603 (0.493) | 0.667 (0.483) | 0.602 (0.492) |
| Range | 0.000 - 1.000 | 0.000 - 1.000 | 0.000 - 1.000 | 0.000 - 1.000 |
| N positive | 16 | 35 | 14 |  |
| **ACR.anti.Smith** |  |  |  |  |
| Mean (SD) | 0.207 (0.412) | 0.276 (0.451) | 0.190 (0.402) | 0.241 (0.430) |
| Range | 0.000 - 1.000 | 0.000 - 1.000 | 0.000 - 1.000 | 0.000 - 1.000 |
| N positive | 6 | 16 | 4 |  |
| **ACR.ANA** |  |  |  |  |
| Mean (SD) | 0.931 (0.258) | 0.966 (0.184) | 0.952 (0.218) | 0.954 (0.211) |
| Range | 0.000 - 1.000 | 0.000 - 1.000 | 0.000 - 1.000 | 0.000 - 1.000 |
| N positive | 27 | 56 | 20 |  |
| **ACR.Hemolytic.Anemia** |  |  |  |  |
| Mean (SD) | 0.138 (0.351) | 0.052 (0.223) | 0.000 (0.000) | 0.065 (0.247) |
| Range | 0.000 - 1.000 | 0.000 - 1.000 | 0.000 - 0.000 | 0.000 - 1.000 |
| N positive | 4 | 3 | 0 |  |
| **ACR.Leukopenia** |  |  |  |  |
| Mean (SD) | 0.138 (0.351) | 0.190 (0.395) | 0.190 (0.402) | 0.176 (0.383) |
| Range | 0.000 - 1.000 | 0.000 - 1.000 | 0.000 - 1.000 | 0.000 - 1.000 |
| N positive | 4 | 11 | 4 |  |
| **ACR.Lymphoenia** |  |  |  |  |
| Mean (SD) | 0.276 (0.455) | 0.293 (0.459) | 0.095 (0.301) | 0.250 (0.435) |
| Range | 0.000 - 1.000 | 0.000 - 1.000 | 0.000 - 1.000 | 0.000 - 1.000 |
| N positive | 8 | 17 | 2 |  |
| **ACR.Thrombocytopenia** |  |  |  |  |
| Mean (SD) | 0.138 (0.351) | 0.190 (0.395) | 0.048 (0.218) | 0.148 (0.357) |
| Range | 0.000 - 1.000 | 0.000 - 1.000 | 0.000 - 1.000 | 0.000 - 1.000 |
| N positive | 4 | 11 | 1 |  |
| **ACR.Renal** |  |  |  |  |
| Mean (SD) | 0.379 (0.494) | 0.500 (0.504) | 0.333 (0.483) | 0.435 (0.498) |
| Range | 0.000 - 1.000 | 0.000 - 1.000 | 0.000 - 1.000 | 0.000 - 1.000 |
| N positive | 11 | 29 | 7 |  |
| **ACR.Photosensitivity** |  |  |  |  |
| Mean (SD) | 0.483 (0.509) | 0.345 (0.479) | 0.429 (0.507) | 0.398 (0.492) |
| Range | 0.000 - 1.000 | 0.000 - 1.000 | 0.000 - 1.000 | 0.000 - 1.000 |
| N positive | 14 | 20 | 9 |  |
| **ACR.APLA** |  |  |  |  |
| Mean (SD) | 0.241 (0.435) | 0.276 (0.451) | 0.286 (0.463) | 0.269 (0.445) |
| Range | 0.000 - 1.000 | 0.000 - 1.000 | 0.000 - 1.000 | 0.000 - 1.000 |
| N positive | 7 | 16 | 6 |  |
| **lupus severity index** |  |  |  |  |
| Mean (SD) | 6.528 (1.670) | 6.886 (1.648) | 6.300 (1.730) | 6.676 (1.672) |
| Range | 3.578 - 9.365 | 3.825 - 9.497 | 3.929 - 8.837 | 3.578 - 9.497 |
| **slicc score** |  |  |  |  |
| Mean (SD) | 1.069 (1.771) | 1.069 (1.425) | 0.667 (0.913) | 0.991 (1.444) |
| Range | 0.000 - 6.000 | 0.000 - 5.000 | 0.000 - 3.000 | 0.000 - 6.000 |
| **sledai score** |  |  |  |  |
| Mean (SD) | 2.069 (2.298) | 2.741 (2.832) | 2.810 (2.639) | 2.574 (2.655) |
| Range | 0.000 - 9.000 | 0.000 - 11.000 | 0.000 - 8.000 | 0.000 - 11.000 |
| **acr lupus nephritis** |  |  |  |  |
| Mean (SD) | 0.379 (0.494) | 0.500 (0.504) | 0.333 (0.483) | 0.435 (0.498) |
| Range | 0.000 - 1.000 | 0.000 - 1.000 | 0.000 - 1.000 | 0.000 - 1.000 |
| **flare severity** |  |  |  |  |
| N-Miss | 18 | 41 | 11 | 70 |
| Mean (SD) | 2.091 (0.701) | 1.412 (0.507) | 2.100 (0.738) | 1.789 (0.704) |
| Range | 1.000 - 3.000 | 1.000 - 2.000 | 1.000 - 3.000 | 1.000 - 3.000 |
| **age at diagnoses** |  |  |  |  |
| Mean (SD) | 25.966 (10.301) | 28.552 (12.031) | 29.810 (11.754) | 28.102 (11.516) |
| Range | 9.000 - 48.000 | 9.000 - 59.000 | 12.000 - 62.000 | 9.000 - 62.000 |

**Table S5: clinical and demographic variables across CD14 clusters**. Flare severity measurement: mild 1, moderate 2, severe 3 . Flare severity measurement: mild 1, moderate 2, severe 3. All are ACR clinical feature are a yes/no variables, except flare, which has 3 levels (mild 1, moderate 2, severe 3). N positive= number of individuals having the clinical feature.

| CD19 | **1 (N=67)** | **2 (N=38)** | **Total (N=105)** |
| --- | --- | --- | --- |
| **ACR.Malar.Rash** |  |  |  |
| Mean (SD) | 0.478 (0.503) | 0.395 (0.495) | 0.448 (0.500) |
| Range | 0.000 - 1.000 | 0.000 - 1.000 | 0.000 - 1.000 |
| N positive | 32 | 15 |  |
| **ACR.Discoid.Rash** |  |  |  |
| Mean (SD) | 0.119 (0.327) | 0.053 (0.226) | 0.095 (0.295) |
| Range | 0.000 - 1.000 | 0.000 - 1.000 | 0.000 - 1.000 |
| N positive | 8 | 2 |  |
| **ACR.Oral.Ulcers** |  |  |  |
| Mean (SD) | 0.537 (0.502) | 0.579 (0.500) | 0.552 (0.500) |
| Range | 0.000 - 1.000 | 0.000 - 1.000 | 0.000 - 1.000 |
| N positive | 36 | 22 |  |
| **ACR.Arthritis** |  |  |  |
| Mean (SD) | 0.791 (0.410) | 0.763 (0.431) | 0.781 (0.416) |
| Range | 0.000 - 1.000 | 0.000 - 1.000 | 0.000 - 1.000 |
| N positive | 53 | 29 |  |
| **ACR.Pleuritis** |  |  |  |
| Mean (SD) | 0.358 (0.483) | 0.237 (0.431) | 0.314 (0.466) |
| Range | 0.000 - 1.000 | 0.000 - 1.000 | 0.000 - 1.000 |
| N positive | 24 | 9 |  |
| **ACR.Pericarditis** |  |  |  |
| Mean (SD) | 0.134 (0.344) | 0.132 (0.343) | 0.133 (0.342) |
| Range | 0.000 - 1.000 | 0.000 - 1.000 | 0.000 - 1.000 |
| N positive | 9 | 5 |  |
| **ACR.Seizure** |  |  |  |
| Mean (SD) | 0.015 (0.122) | 0.132 (0.343) | 0.057 (0.233) |
| Range | 0.000 - 1.000 | 0.000 - 1.000 | 0.000 - 1.000 |
| N positive | 1 | 5 |  |
| **ACR.Psychosis** |  |  |  |
| Mean (SD) | 0.060 (0.239) | 0.026 (0.162) | 0.048 (0.214) |
| Range | 0.000 - 1.000 | 0.000 - 1.000 | 0.000 - 1.000 |
| N positive | 4 | 1 |  |
| **ACR.anti.dsDNA** |  |  |  |
| Mean (SD) | 0.597 (0.494) | 0.605 (0.495) | 0.600 (0.492) |
| Range | 0.000 - 1.000 | 0.000 - 1.000 | 0.000 - 1.000 |
| N positive | 40 | 23 |  |
| **ACR.anti.Smith** |  |  |  |
| Mean (SD) | 0.269 (0.447) | 0.184 (0.393) | 0.238 (0.428) |
| Range | 0.000 - 1.000 | 0.000 - 1.000 | 0.000 - 1.000 |
| N positive | 18 | 7 |  |
| **ACR.ANA** |  |  |  |
| Mean (SD) | 0.970 (0.171) | 0.921 (0.273) | 0.952 (0.214) |
| Range | 0.000 - 1.000 | 0.000 - 1.000 | 0.000 - 1.000 |
| N positive | 65 | 35 |  |
| **ACR.Hemolytic.Anemia** |  |  |  |
| Mean (SD) | 0.030 (0.171) | 0.105 (0.311) | 0.057 (0.233) |
| Range | 0.000 - 1.000 | 0.000 - 1.000 | 0.000 - 1.000 |
| N positive | 2 | 4 |  |
| **ACR.Leukopenia** |  |  |  |
| Mean (SD) | 0.194 (0.398) | 0.211 (0.413) | 0.200 (0.402) |
| Range | 0.000 - 1.000 | 0.000 - 1.000 | 0.000 - 1.000 |
| N positive | 13 | 8 |  |
| **ACR.Lymphoenia** |  |  |  |
| Mean (SD) | 0.224 (0.420) | 0.395 (0.495) | 0.286 (0.454) |
| Range | 0.000 - 1.000 | 0.000 - 1.000 | 0.000 - 1.000 |
| N positive | 15 | 15 |  |
| **ACR.Thrombocytopenia** |  |  |  |
| Mean (SD) | 0.119 (0.327) | 0.184 (0.393) | 0.143 (0.352) |
| Range | 0.000 - 1.000 | 0.000 - 1.000 | 0.000 - 1.000 |
| N positive | 8 | 7 |  |
| **ACR.Renal** |  |  |  |
| Mean (SD) | 0.403 (0.494) | 0.526 (0.506) | 0.448 (0.500) |
| Range | 0.000 - 1.000 | 0.000 - 1.000 | 0.000 - 1.000 |
| N positive | 27 | 20 |  |
| **ACR.Photosensitivity** |  |  |  |
| Mean (SD) | 0.388 (0.491) | 0.316 (0.471) | 0.362 (0.483) |
| Range | 0.000 - 1.000 | 0.000 - 1.000 | 0.000 - 1.000 |
| N positive | 26 | 12 |  |
| **ACR.APLA** |  |  |  |
| Mean (SD) | 0.239 (0.430) | 0.395 (0.495) | 0.295 (0.458) |
| Range | 0.000 - 1.000 | 0.000 - 1.000 | 0.000 - 1.000 |
| N positive | 16 | 15 |  |
| **lupus severity index** |  |  |  |
| Mean (SD) | 6.475 (1.712) | 7.174 (1.484) | 6.728 (1.660) |
| Range | 3.825 - 9.047 | 3.578 - 9.497 | 3.578 - 9.497 |
| **slicc score** |  |  |  |
| Mean (SD) | 0.836 (1.214) | 1.474 (1.842) | 1.067 (1.495) |
| Range | 0.000 - 5.000 | 0.000 - 6.000 | 0.000 - 6.000 |
| **sledai score** |  |  |  |
| Mean (SD) | 2.836 (3.208) | 2.395 (2.574) | 2.676 (2.989) |
| Range | 0.000 - 16.000 | 0.000 - 9.000 | 0.000 - 16.000 |
| **acr lupus nephritis** |  |  |  |
| Mean (SD) | 0.403 (0.494) | 0.500 (0.507) | 0.438 (0.499) |
| Range | 0.000 - 1.000 | 0.000 - 1.000 | 0.000 - 1.000 |
| **flare severity** |  |  |  |
| N-Miss | 41 | 29 | 70 |
| Mean (SD) | 1.846 (0.784) | 1.778 (0.441) | 1.829 (0.707) |
| Range | 1.000 - 3.000 | 1.000 - 2.000 | 1.000 - 3.000 |
| **age at diagnoses** |  |  |  |
| Mean (SD) | 27.313 (10.734) | 31.395 (14.200) | 28.790 (12.196) |
| Range | 9.000 - 57.000 | 9.000 - 64.000 | 9.000 - 64.000 |

**Table S6: clinical and demographic variables across CD19 clusters.** Flare severity measurement: mild 1, moderate 2, severe 3 . Flare severity measurement: mild 1, moderate 2, severe 3. All are ACR clinical feature are a yes/no variables, except flare, which has 3 levels (mild 1, moderate 2, severe 3). N positive= number of individuals having the clinical feature.

| NK | **1 (N=79)** | **2 (N=12)** | **Total (N=91)** |
| --- | --- | --- | --- |
| **ACR.Malar.Rash** |  |  |  |
| Mean (SD) | 0.430 (0.498) | 0.333 (0.492) | 0.418 (0.496) |
| Range | 0.000 - 1.000 | 0.000 - 1.000 | 0.000 - 1.000 |
| N positive | 34 | 4 |  |
| **ACR.Discoid.Rash** |  |  |  |
| Mean (SD) | 0.089 (0.286) | 0.250 (0.452) | 0.110 (0.314) |
| Range | 0.000 - 1.000 | 0.000 - 1.000 | 0.000 - 1.000 |
| N positive | 7 | 3 |  |
| **ACR.Oral.Ulcers** |  |  |  |
| Mean (SD) | 0.544 (0.501) | 0.833 (0.389) | 0.582 (0.496) |
| Range | 0.000 - 1.000 | 0.000 - 1.000 | 0.000 - 1.000 |
| N positive | 43 | 10 |  |
| **ACR.Arthritis** |  |  |  |
| Mean (SD) | 0.747 (0.438) | 0.917 (0.289) | 0.769 (0.424) |
| Range | 0.000 - 1.000 | 0.000 - 1.000 | 0.000 - 1.000 |
| N positive | 59 | 11 |  |
| **ACR.Pleuritis** |  |  |  |
| Mean (SD) | 0.304 (0.463) | 0.417 (0.515) | 0.319 (0.469) |
| Range | 0.000 - 1.000 | 0.000 - 1.000 | 0.000 - 1.000 |
| N positive | 24 | 5 |  |
| **ACR.Pericarditis** |  |  |  |
| Mean (SD) | 0.165 (0.373) | 0.083 (0.289) | 0.154 (0.363) |
| Range | 0.000 - 1.000 | 0.000 - 1.000 | 0.000 - 1.000 |
| N positive | 13 | 1 |  |
| **ACR.Seizure** |  |  |  |
| Mean (SD) | 0.076 (0.267) | 0.000 (0.000) | 0.066 (0.250) |
| Range | 0.000 - 1.000 | 0.000 - 0.000 | 0.000 - 1.000 |
| N positive | 6 | 0 |  |
| **ACR.Psychosis** |  |  |  |
| Mean (SD) | 0.038 (0.192) | 0.083 (0.289) | 0.044 (0.206) |
| Range | 0.000 - 1.000 | 0.000 - 1.000 | 0.000 - 1.000 |
| N positive | 3 | 1 |  |
| **ACR.anti.dsDNA** |  |  |  |
| Mean (SD) | 0.570 (0.498) | 0.583 (0.515) | 0.571 (0.498) |
| Range | 0.000 - 1.000 | 0.000 - 1.000 | 0.000 - 1.000 |
| N positive | 45 | 7 |  |
| **ACR.anti.Smith** |  |  |  |
| Mean (SD) | 0.278 (0.451) | 0.333 (0.492) | 0.286 (0.454) |
| Range | 0.000 - 1.000 | 0.000 - 1.000 | 0.000 - 1.000 |
| N positive | 22 | 4 |  |
| **ACR.ANA** |  |  |  |
| Mean (SD) | 0.962 (0.192) | 0.833 (0.389) | 0.945 (0.229) |
| Range | 0.000 - 1.000 | 0.000 - 1.000 | 0.000 - 1.000 |
| N positive | 76 | 10 |  |
| **ACR.Hemolytic.Anemia** |  |  |  |
| Mean (SD) | 0.076 (0.267) | 0.000 (0.000) | 0.066 (0.250) |
| Range | 0.000 - 1.000 | 0.000 - 0.000 | 0.000 - 1.000 |
| N positive | 6 | 0 |  |
| **ACR.Leukopenia** |  |  |  |
| Mean (SD) | 0.203 (0.404) | 0.167 (0.389) | 0.198 (0.401) |
| Range | 0.000 - 1.000 | 0.000 - 1.000 | 0.000 - 1.000 |
| N positive | 16 | 2 |  |
| **ACR.Lymphoenia** |  |  |  |
| Mean (SD) | 0.266 (0.445) | 0.167 (0.389) | 0.253 (0.437) |
| Range | 0.000 - 1.000 | 0.000 - 1.000 | 0.000 - 1.000 |
| N positive | 21 | 2 |  |
| **ACR.Thrombocytopenia** |  |  |  |
| Mean (SD) | 0.152 (0.361) | 0.083 (0.289) | 0.143 (0.352) |
| Range | 0.000 - 1.000 | 0.000 - 1.000 | 0.000 - 1.000 |
| N positive | 12 | 1 |  |
| **ACR.Renal** |  |  |  |
| Mean (SD) | 0.443 (0.500) | 0.333 (0.492) | 0.429 (0.498) |
| Range | 0.000 - 1.000 | 0.000 - 1.000 | 0.000 - 1.000 |
| N positive | 35 | 4 |  |
| **ACR.Photosensitivity** |  |  |  |
| Mean (SD) | 0.367 (0.485) | 0.583 (0.515) | 0.396 (0.492) |
| Range | 0.000 - 1.000 | 0.000 - 1.000 | 0.000 - 1.000 |
| N positive | 29 | 7 |  |
| **ACR.APLA** |  |  |  |
| Mean (SD) | 0.304 (0.463) | 0.167 (0.389) | 0.286 (0.454) |
| Range | 0.000 - 1.000 | 0.000 - 1.000 | 0.000 - 1.000 |
| N positive | 24 | 2 |  |
| **lupus severity index** |  |  |  |
| Mean (SD) | 6.758 (1.720) | 6.084 (1.686) | 6.670 (1.722) |
| Range | 3.578 - 9.497 | 4.088 - 8.336 | 3.578 - 9.497 |
| **slicc score** |  |  |  |
| Mean (SD) | 1.127 (1.612) | 0.417 (0.669) | 1.033 (1.538) |
| Range | 0.000 - 6.000 | 0.000 - 2.000 | 0.000 - 6.000 |
| **sledai score** |  |  |  |
| Mean (SD) | 2.886 (3.158) | 2.000 (1.651) | 2.769 (3.011) |
| Range | 0.000 - 16.000 | 0.000 - 6.000 | 0.000 - 16.000 |
| **acr lupus nephritis** |  |  |  |
| Mean (SD) | 0.443 (0.500) | 0.333 (0.492) | 0.429 (0.498) |
| Range | 0.000 - 1.000 | 0.000 - 1.000 | 0.000 - 1.000 |
| **flare severity** |  |  |  |
| N-Miss | 54 | 4 | 58 |
| Mean (SD) | 1.920 (0.640) | 1.625 (0.744) | 1.848 (0.667) |
| Range | 1.000 - 3.000 | 1.000 - 3.000 | 1.000 - 3.000 |
| **age at diagnoses** |  |  |  |
| Mean (SD) | 27.557 (12.050) | 32.000 (10.617) | 28.143 (11.912) |
| Range | 9.000 - 64.000 | 14.000 - 48.000 | 9.000 - 64.000 |

**Table S7: clinical and demographic variables across NK clusters** Flare severity measurement: mild 1, moderate 2, severe 3. Flare severity measurement: mild 1, moderate 2, severe 3. All are ACR clinical feature are a yes/no variables, except flare, which has 3 levels (mild 1, moderate 2, severe 3). N positive= number of individuals having the clinical feature.

|  | **Lupus severity index** | **SLICC score** | **SLEDAI score** | **Race**  **(Whites vs Asians)** | **Lupus nephritis** | **Age at diagnoses** |
| --- | --- | --- | --- | --- | --- | --- |
| CD4 | 6 | 1 | 0 | 11 | 3 | 0 |
| CD14 | 1 | 0 | 0 | 14 | 2 | 1 |
| CD19 | 3 | 2 | 2 | 18 | 2 | 0 |
| NK | 29 | 0 | 3 | 56 | 2 | 0 |

**Table S8:** Number of statistically significant genes (p value adjusted < 0.05 & abs (log2FC) 1) for differential expression analyses conducted on lupus severity score, SLICC score, SLEDAI score, Race (White vs Asians), Lupus nephritis, and Age at diagnoses (late vs early onset).

**Supplementary Figures:**

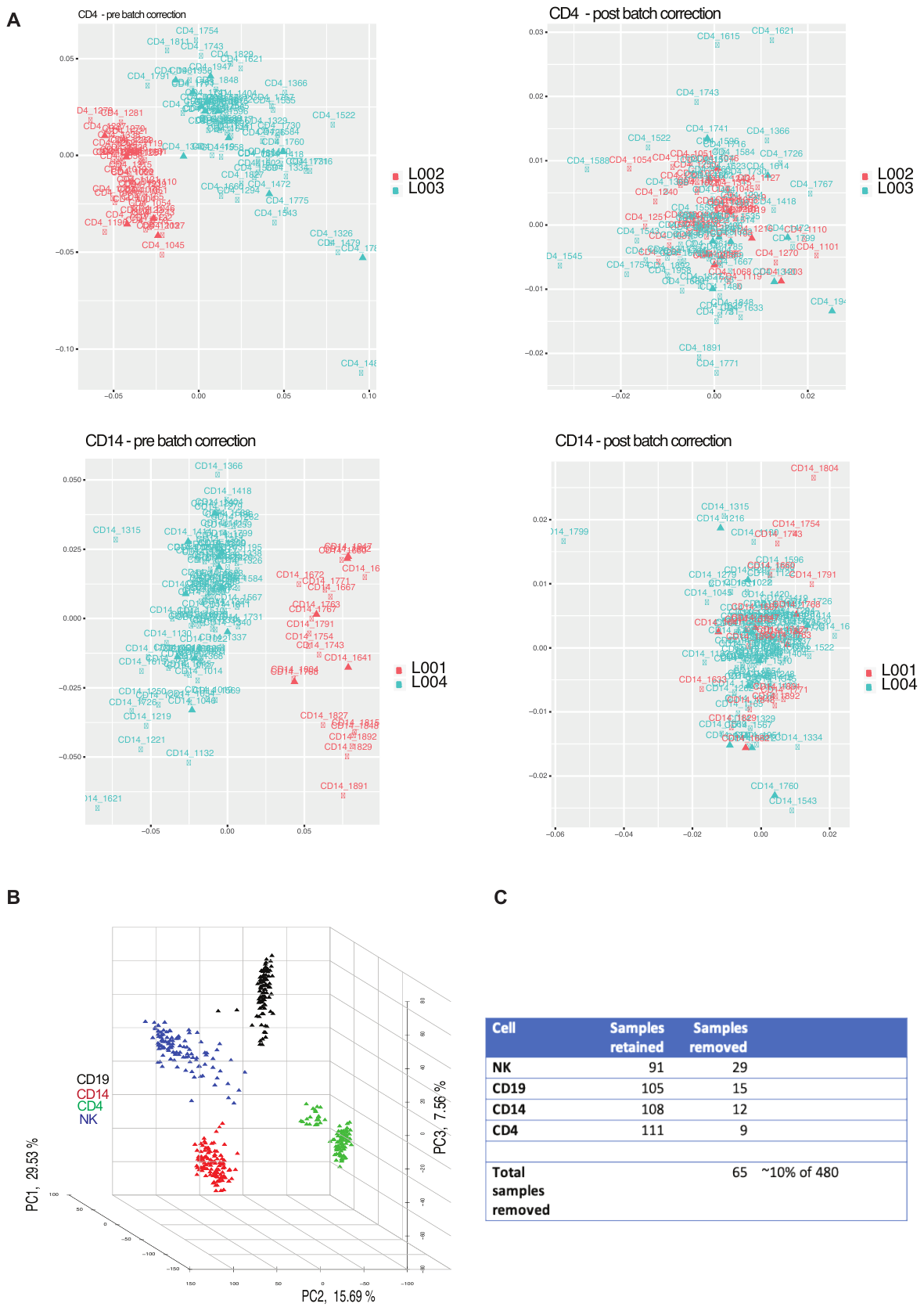

**Figure S1:** A) Data before and after batch correction using limma. Batch effect was observed only in CD4^+^ T cells and CD14^+^ monocytes. K-means clustering after batch correction. B) Principal component analysis (PCA) on the batch corrected data. C) Table detailing the number of samples removed per immune-cell type after QC.

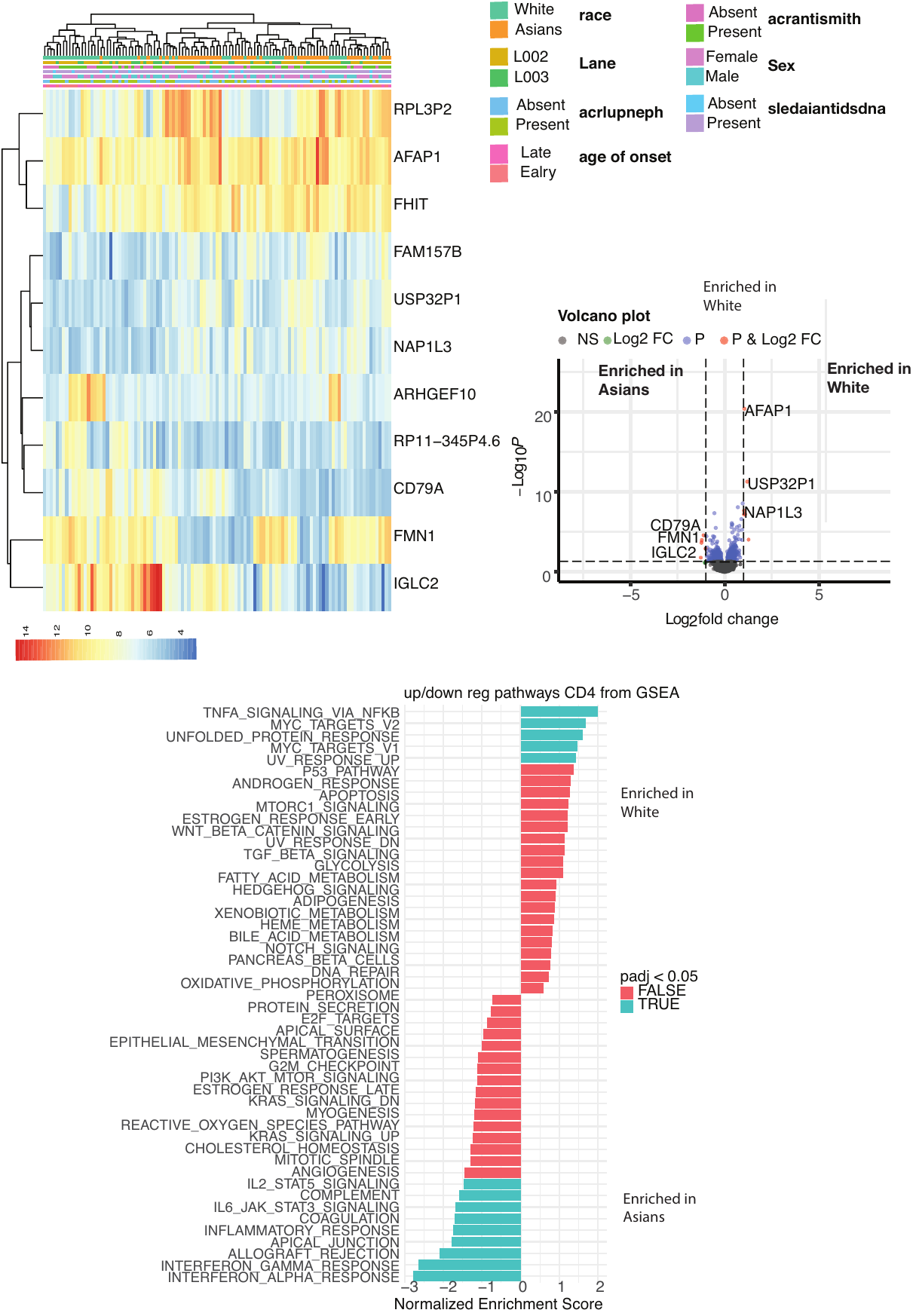

**Figure S2:** Differential Expression by race for CD4^+^ T cell.

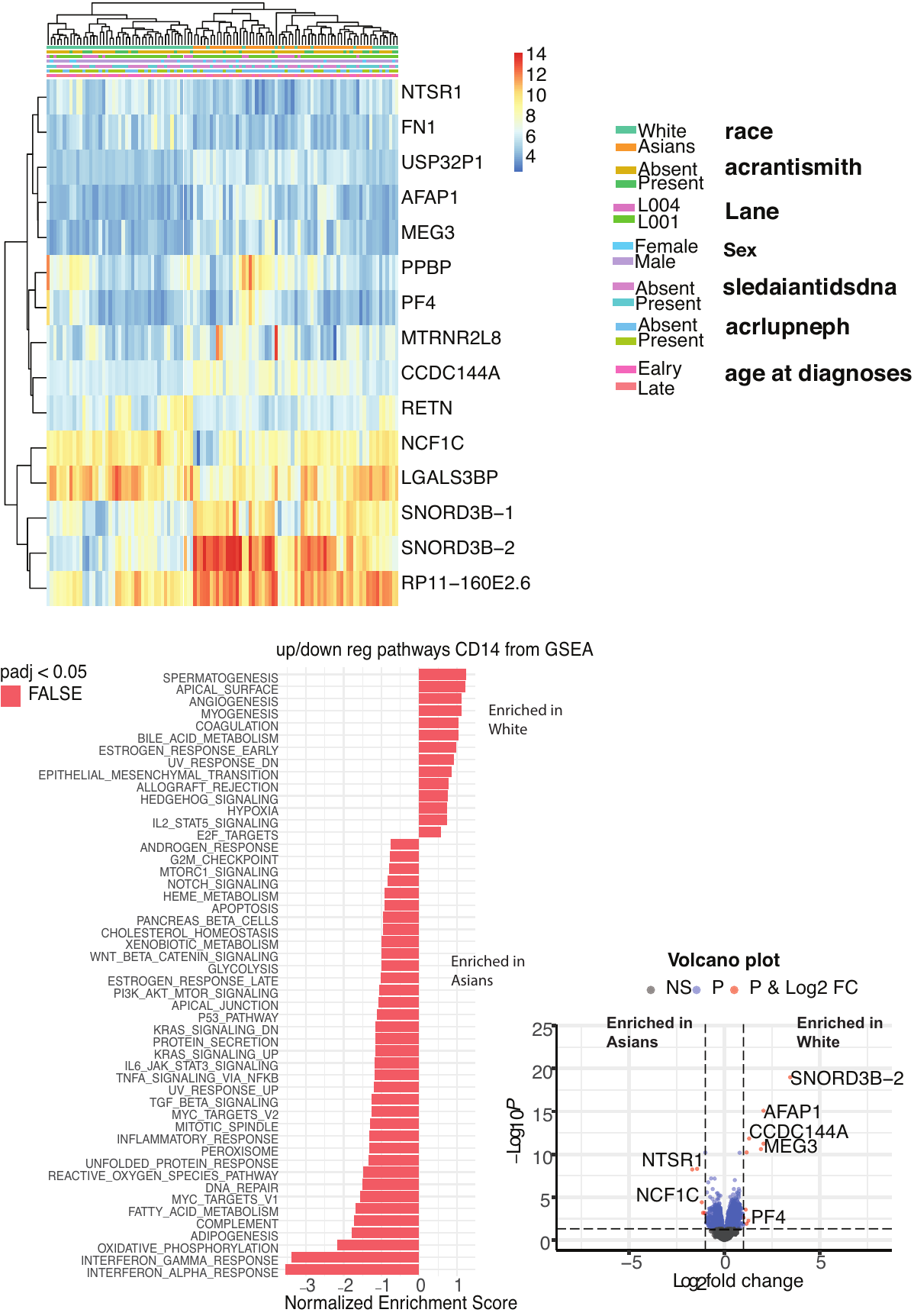
**Figure S3:** Differential Expression by race for CD14^+^ monocytes.

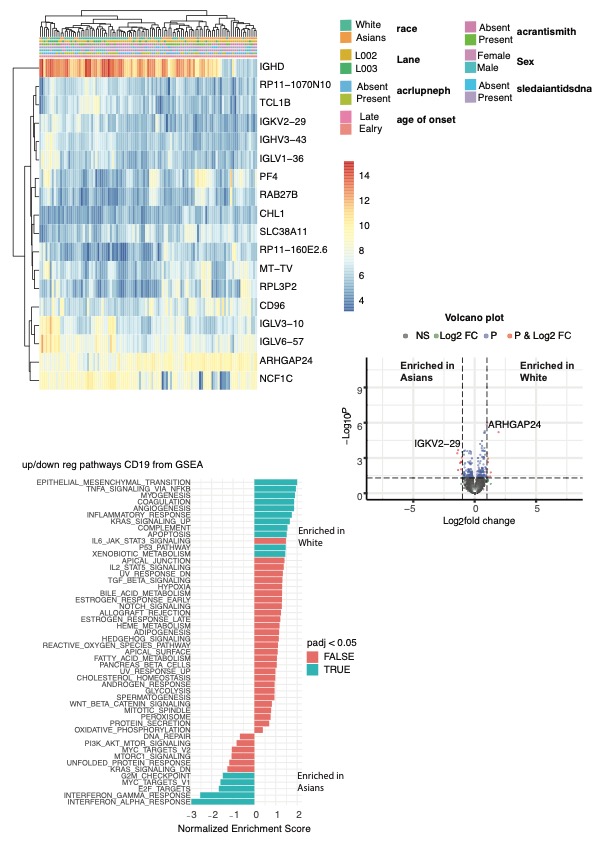
**Figure S4:** Differential Expression by race for CD19 B cell.

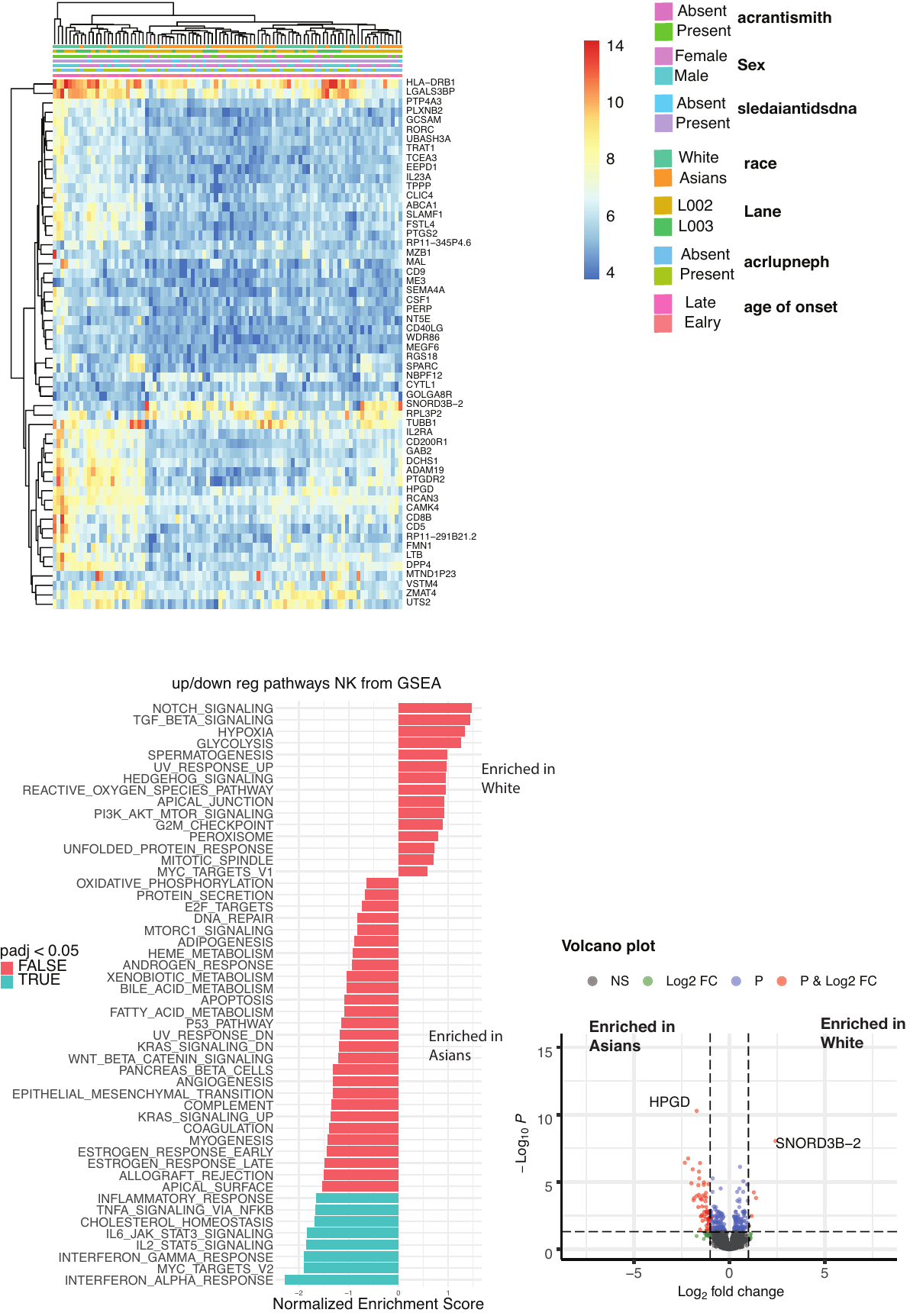
**Figure S5:** Differential Expression by race for NK cells.

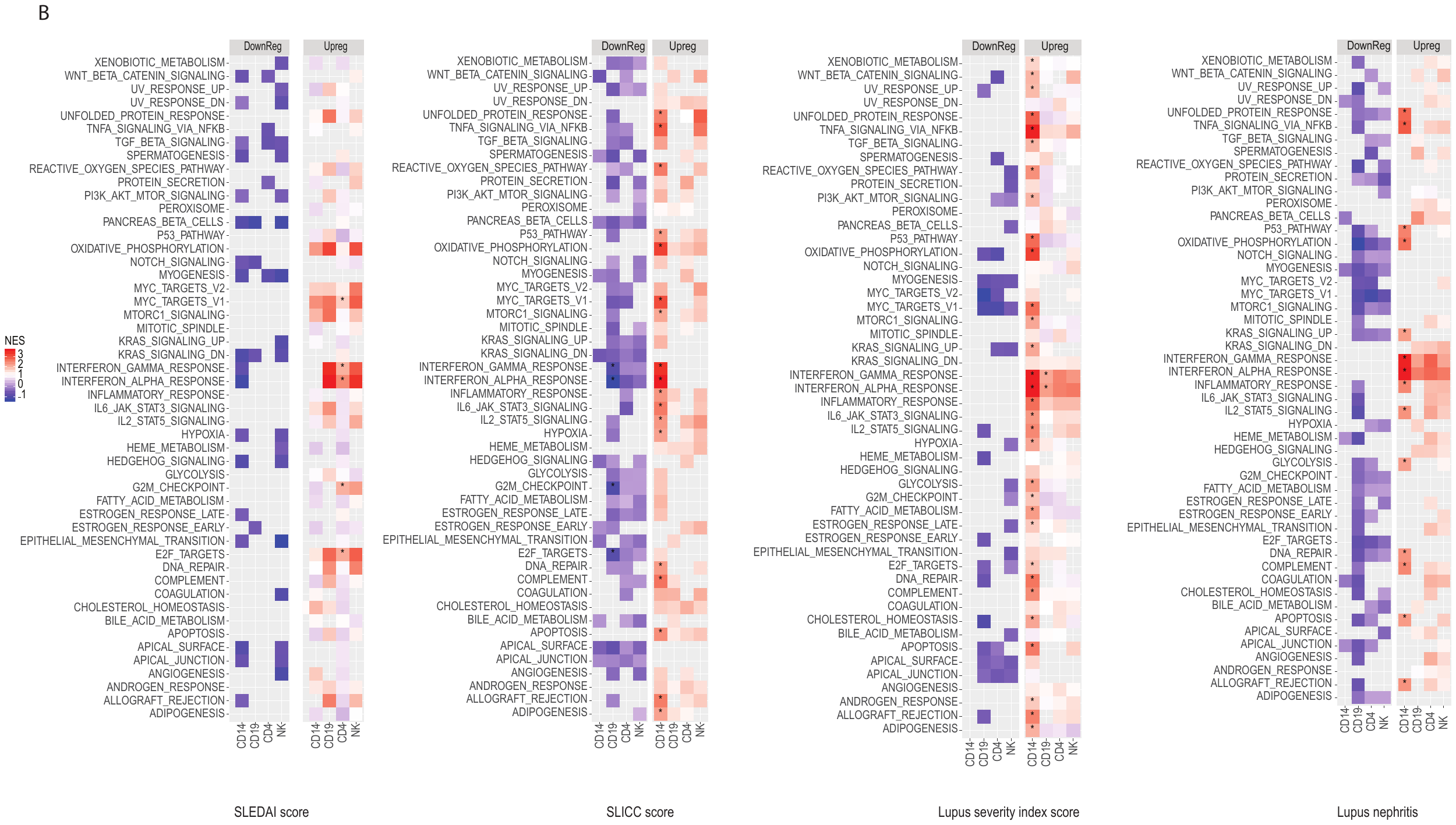

**Figure S6:** Pathway Analysis using GSEA from DE conducted on other clinical criteria Sledai score, slicc score, lupus severity index, and lupus nephritis

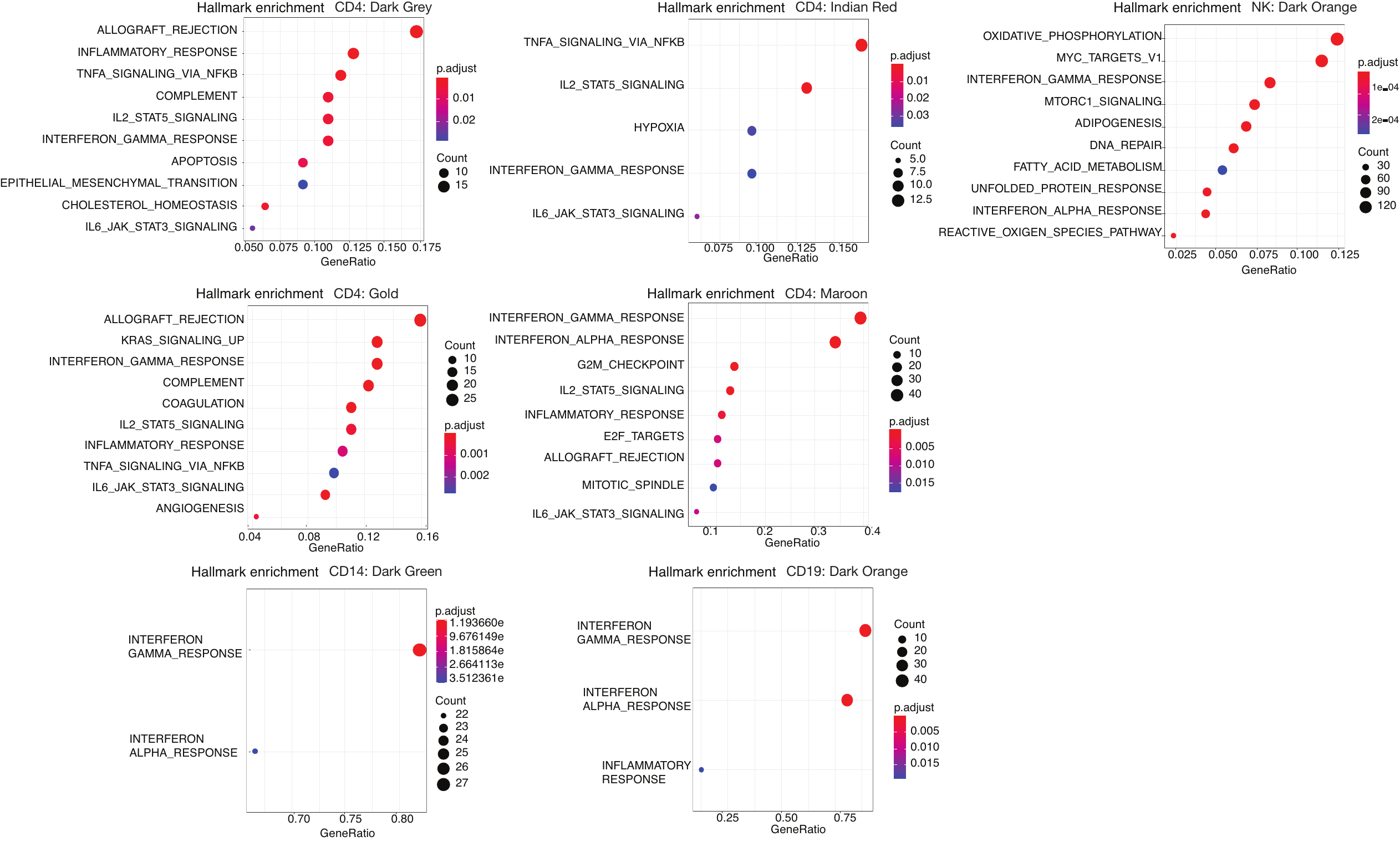

**Figure S7:** Functional enrichment (Hallmark pathways) of co-expressed modules enriched for interferon pathways.

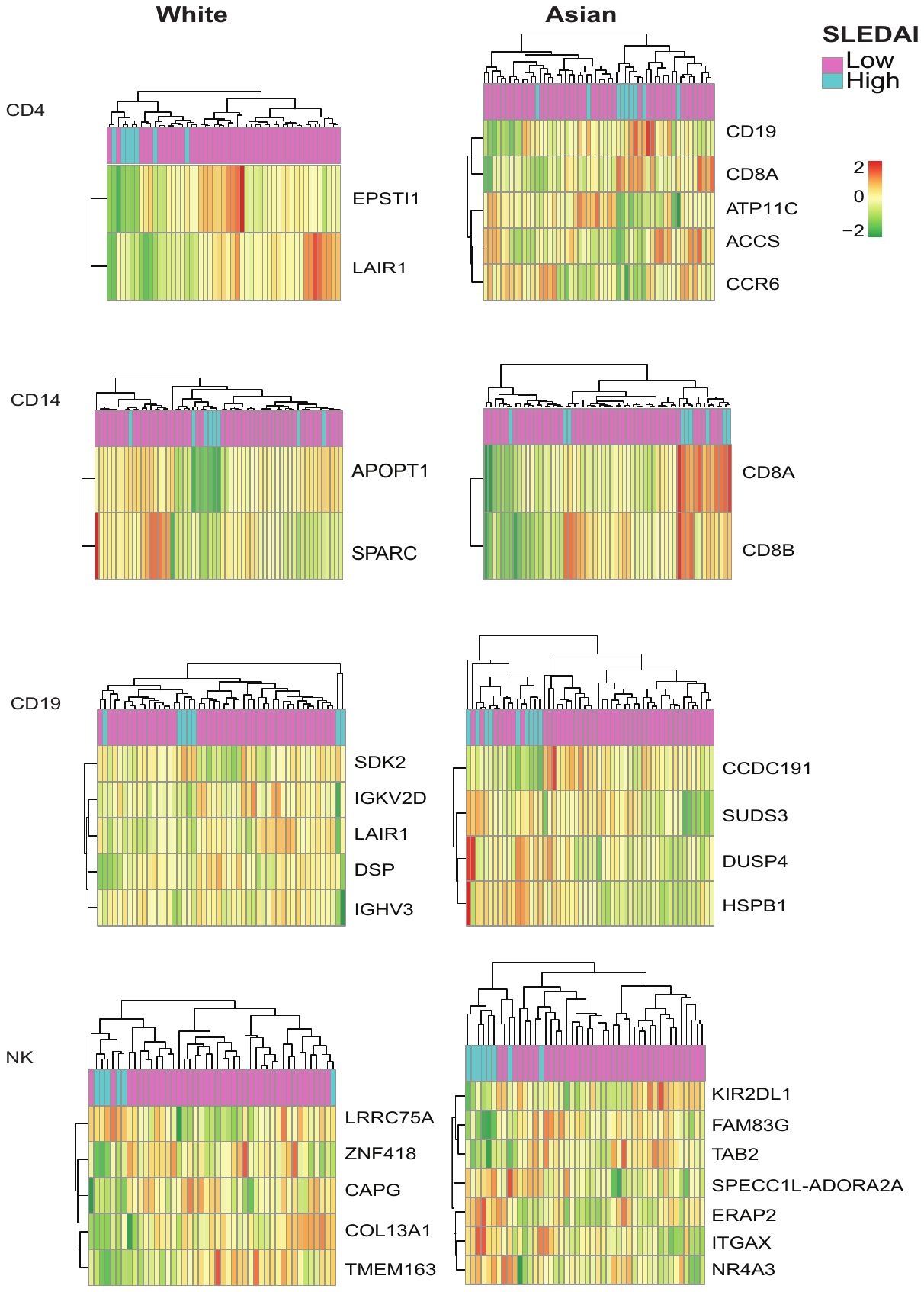

**Figure S8:** Clustergrams by cell type for each ethnic group by using the expression values of the top predictors.
